# Experimental validation of genome-environment associations in Arabidopsis

**DOI:** 10.1101/2025.01.08.631904

**Authors:** Yuxin Luo, Claire M. Lorts, Erica H. Lawrence-Paul, Jesse R. Lasky

## Abstract

Identifying the genetic basis of local adaptation is a key goal in evolutionary biology. Allele frequency clines along environmental gradients, known as genotype-environment associations (GEA), are often used to detect potential loci causing local adaptation but are rarely followed by experimental validation. Here, we tested loci identified in three moisture-related GEA studies on *Arabidopsis*. We studied 42 GEA-identified genes using t-DNA knockout lines under drought and tested effects on flowering time, an adaptive trait, and genotype-by-environment (GxE) interactions on performance and fitness. In total, 16/42 genes had significant effects on traits involved in local adaptation or performance response to environment. We found that *wrky38* mutants had significant GxE effects for fitness; *lsd1* plants had a significant GxE effect for flowering time, while 11 genes showed flowering time effects with no drought interaction. However, most GEA candidates did not exhibit GxE. In the follow-up experiments, *wrky38* caused decreased stomatal conductance and specific leaf area under drought, indicating potentially adaptive drought avoidance. Additionally, GEA identified natural putative LoF variants of *WRKY38* associated with dry environments, as well as alleles associated with variation in *LSD1* expression. While only a few GEA-identified genes were validated for GxE interactions for fitness under drought, we likely overlooked some because experiments might not well represent natural environments and t-DNA insertions might not well represent natural alleles. Nevertheless, GEAs apparently identified some genes contributing to local adaptation. GEA and follow-up experiments are straightforward to implement in model systems and demonstrate prospects for GEA discovery of new local adaptations.

## INTRODUCTION

A substantial portion of genetic variation within species in ecologically important traits is likely due to local adaptation to the environment (Savolainen et al., 2013; Tigano and Friesen, 2016; Wadgymar et al., 2022). Local adaptation is defined by a genotype-by-environment (GxE) interaction where local genotypes have higher fitness than foreign genotypes (Kawecki & Ebert, 2004). Identifying locally adaptive alleles and traits is of great interest in plant and animal breeding and conservation, where these alleles can be deployed for genetic improvement and prediction (Cortés et al., 2022; Crossa et al., 2017; Lasky et al., 2023; Whiting et al., 2024). Central questions of local adaptation include the effect size and number of involved mutations (Yeaman and Whitlock, 2011), whether locally adapted mutations have tradeoffs among different environments (Lee et al., 2024), whether the same or different mutations are involved in local adaptation to the same environments in different populations (Ralph and Coop, 2015), and which environmental gradients drive locally adaptive genetic variation (Lasky et al., 2012).

Conducting multiple common garden experiments to map populations and genetic loci contributing to fitness tradeoffs has been a gold standard approach (Clausen et al., 1941). However, such experiments are logistically challenging or impossible in many systems. Alternative approaches using geographic patterns in allele frequency have emerged for identifying loci potentially underlying local adaptation. Loci showing substantial allele frequency differences between populations, i.e. certain environment variables, may indicate those driving local adaptation (Coop et al., 2010; Endler, 1973). With the increasing availability of high-throughput genomic data, Genome-Environment Association (GEA) studies have been widely employed to identify putative locally adapted loci (Lasky et al., 2023; Rellstab et al., 2015).

GEAs employ diverse methods to identify loci where allele frequency is closely associated with environmental conditions (Lasky et al., 2023; Rellstab et al., 2015). Methods differ depending on whether they are single-locus or multi-loci (e.g. genome-wide) models (Betancourt et al., 2021; Gehan et al., 2015; Hager et al., 2022; Hancock et al., 2011; Lasky et al., 2012; Lee et al., 2024), where the latter may capture groups of loci covarying with the environment that are potentially relevant for polygenic adaptation (Forester et al., 2018). Additionally, there are methods that control for genome-wide patterns of similarity among populations using random effects (Zoubarev et al., 2012). While these approaches may reduce spurious associations due to population structure-environmental covariation, they can also be too conservative (Alonso-Blanco et al., 2016). Thirdly, some approaches synthesize the evidence from GEA with additional evidence, such as multiple common garden experiments (Capblancq et al., 2023; Lasky et al., 2018; Soudi et al., 2023). While these methods are now standard approaches in ecological genomics, the findings are rarely followed by experimental validation of variation at individual loci (Monroe et al., 2018; Nyine et al., 2025; Tergemina et al., 2025).

To experimentally test the hypothesis that variation at a given locus causes GxE for fitness, approaches that isolate experimental genetic variation to that locus are the most powerful. Creating near-isogenic lines (NILs) segregating at a given locus is ideal for testing GxE effects in a natural genetic background, but it is a time-consuming process. Gene knockout mutants in model species like *Arabidopsis* provide an alternative tool for testing GxE effects of variation at a particular gene. For example, various Arabidopsis knockout lines carrying t-DNA insertions within genes that disrupt their function provide valuable resources for studying the effects of individual genes on fitness and other ecologically important traits. Although these mutants may not fully represent natural allelic variation for local adaptation, using them may be a good option for bulk experimental screening of candidate loci (Chong and Stinchcombe, 2019; Monroe et al., 2018). Resequencing of large numbers of natural *Arabidopsis* genotypes has yielded more detailed information on functional variation (Alonso-Blanco et al., 2016), which can be linked to loci of interest to make inferences about functional variants generating strong signals in GEA studies.

*Arabidopsis thaliana* is a small annual plant native to a wide range across Eurasia and Africa with diverse climates (Yim et al., 2024). Many studies have demonstrated potential drought adaptation of *A. thaliana* in quantitative traits (Dittberner et al., 2018), SNPs or genes (El-Soda et al., 2015; Exposito-Alonso et al., 2018; Hancock et al., 2011), and gene expression (Lasky et al., 2014). *A. thaliana* employs diverse strategies for drought adaptation such as drought escape via phenology, drought avoidance via traits reducing water loss, and drought tolerance via enhancing water acquisition or water-use efficiency (Lovell et al., 2013).

Here we focus on studying loci identified based on 3 different approaches, with the goal of testing whether identified genes indeed cause genotype-drought effects on fitness or other aspects of performance. We are also interested in whether the genes identified as drought-adapted in our experiments have allele frequency clines across the worldwide distribution of *A. thaliana*. For mutants identified as showing interesting genotype-environment interactions in our initial drought screen experiment, we followed up by testing their response to an additional, different drought treatment.

We tested the following hypotheses:

1. Some genes identified through GEA as potentially drought adapted exhibit natural variation in putative gene function assessed from resequencing data. Such genes would be especially amenable to using t-DNA knockout lines for testing the effects of natural variation. Mutations in genes without natural loss-of-function variation can still be assessed using t-DNA lines, though their relevance to natural variants is less clear.
2. Knockout mutants of these genes display significant genotype-drought interactions affecting traits (e.g. flowering time) known to be under changing selection in drought, fitness or performance traits in controlled drought experiments; and
3. These genes show climate-associated allelic variation and genomic signatures of selection, patterns consistent with adaptation to dry climates.

In our main drought screen, we found that *wrky38* had significant GxE effects on several fitness-related traits, and *lsd1* had a significant GxE effect on flowering time. Both *WRKY38* and *LSD1* were top candidates associated with moisture in their originated studies (Lasky et al., 2018, 2012). Additionally, *WRKY38* was also found most strongly associated with winter low temperatures in the combined common garden study (Lasky et al., 2018), while knockouts of *LSD1* were found to affect chilling sensitivity (Huang et al., 2010) and allelic variation was associated with the second principal component of climate (moisture related, Lee et al., 2017) in Arabidopsis. Both genes happen to play a role in the salicylic acid (SA) pathway (Bernacki et al., 2019; Kim et al., 2008; Szechyńska-Hebda et al., 2016; Wituszynska et al., 2013), primarily known for microbial defense response but which can be activated by abiotic stress (reviewed in Miura and Tada, 2014; Wu et al., 2019). Incorporating previous studies and our results, we decided to use the two mutants to follow up with intermediate drought and freezing experiments, in order to test their possible adaptation in different drought and cold regimes in terms of their ecophysiological responses.

## METHODS

### Selecting genes from published genome-environment associations

We selected genes to test using five SNP lists from three published GEA studies that employed three different approaches. The first approach was based on mixed models of genome-wide associations with climate, where we included random effects to account for genomic similarity (Kang et al., 2008). For these mixed models, we calculated SNP associations with intra-annual monthly precipitation variability, growing season precipitation variability, or inter-annual growing season precipitation variability (i.e. 3 separate environmental GWAS, Lasky et al., 2014). We took the SNPs from all three climate associations and combined them into one list ranked by *p*-value for the mixed model association test. The second approach was a multivariate ordination where we identified SNPs most strongly associated with the first axis, which was associated with seasonality of temperature and precipitation, or the first axis after partialing out spatial variables, which was strongly associated with summer moisture (Lasky et al., 2012). We took SNPs with the strongest absolute loadings on each of these axes. The third approach was based on integrated information from four common gardens across Europe and also the broader ecotype panel to identify SNPs that showed the strongest SNP-climate associations and GxE for fitness favoring home climate alleles in monthly growing season precipitation variability or aridity index (i.e. annual precipitation/PET; Lasky et al., 2018). In total, we generated 1, 2, and 2 lists of SNPs from the three studies respectively.

From the top SNPs in each list, we only kept SNPs at least 250 kb apart. Starting from the SNP with the lowest *p*-values, we took up to 3 closest genes within 5kb of the SNP until we had at least 15 genes for each of the four lists of SNPs, or in the case of the combined climate association study, until we had at least 25 genes.

This procedure resulted in a list of 90 candidate genes. We chose 44 SALK lines representing knockouts of 42 genes, including 42 lines with insertions within exons and 2 lines with insertions within introns and confirmed homozygosity using tDNA Express: Arabidopsis Gene Mapping Tool (Alonso et al., 2003; Table S1). Among the 44 SALK lines targeting 42 genes, three lines (SALK_048316C, SALK_085852C, and SALK_063969) targeting AT5G65080 (*MAF5*) and AT3G45860 (*CRK4*), although not top candidates, were included in the experiment because their potential functional significance in drought adaptation (Kim and Sung, 2010; Ratcliffe et al., 2003; Zhao et al., 2023). As a result, 40 top-candidate genes were validated in this study.

### Functional variation in the screened loci in the 1001 Genomes

We analyzed putative functional variation (hereafter FV) in the 1001 Genomes resequencing data for all genes screened. We used the 1001 Genomes Polymorphism Browser (https://tools.1001genomes.org/polymorph/) to identify potential large-effect mutations in each gene, which were considered as putative loss-of-function (LoF) mutations. If more than one isoform existed for a gene, we recorded the case of the most common isoform, except when no functional variant was found for this isoform, we instead used the most common isoform that had functional variants. We summarized the counts of each type of functional variation and the positions where the variations occurred.

### Main drought screen

Detailed experimental settings are provided in Supplementary Methods. Briefly, we programmed the chamber temperature and photoperiod based on that of the *Lip-0* ecotype from southern Poland following Lorts and Lasky (2020) (Table S2). Each tray represented either a drought or well-watered treatment, and seven replicates per mutant were randomly assigned positions for each treatment. Seeds were stratified in DI water for 4-5 days prior to planting.

Pots were maintained at 2.5-5 cm of bottom water before drought treatments, which began 19 days post-planting. Bottom water was removed from drought trays, while well-watered trays retained a constant 2.5 cm of water and received 2.5 ml of top water every other day. After five weeks, all pots were top-watered with a 12 ml miracle grow solution every other day to ensure nutrition.

At maturity, fecundity was assessed when siliques dried and rosette leaves senesced. We recorded inflorescence height and secondary branch numbers were recorded. For reproductive fitness estimates, we measured total silique number and silique length for 6 siliques to calculate average silique length, then calculated the total silique length as the product of silique number and average silique length.

After measurements, aboveground dry biomass was obtained, and rosette tissue was ground for δ^13^C and δ^15^N isotope analysis. δ^13^C represents estimate for water use efficiency (WUE).

### Follow-up drought and freezing experiments

#### Intermediate drought experiment on *wrky38* and *lsd1*

For intermediate drought (ID), plants were grown at 20/14°C (12h/12h) with varying watering frequencies (every 3 days for well-watered, every 6 days for drought). We tracked stomatal conductance (g_sw_) and Fv/Fm for 10 for four plants per genotype per treatment. Specific leaf area (SLA), relative water content (RWC), and leaf isotope analyses (δ^13^C, C:N and δ^15^N) were analyzed using five to six 40-day-old rosettes (see Supplementary Methods for details).

We recorded the flowering time for six replicates per genotype per treatment. As in the main drought screen, we measured the same growth and fitness traits and isotopes.

#### Overnight freezing experiment on *wrky38* and *lsd1*

Twelve replicates of *Col*, *wrky38*, and *lsd1* were grown at 20/14°C (12h/12h) until Day 30, then acclimated to 10/4℃ for 2 weeks before freezing treatment at 10/-2℃. We surrounded all pots with extra soil for insulation to prevent unnaturally soil freezing. On day 60, we measured diameter as an estimate of growth under freezing. After temperature was restored on Day 64, flowering time, biomass, silique traits, and inflorescence length were recorded as described above.

### Statistical analyses

In both the drought experiments, we were primarily interested in whether the knockouts had genotype-specific drought responses, i.e., whether there were genotype-by-drought interactions, as well as whether the knockouts altered average phenotypes under drought, i.e., whether genotype effects were significant. Thus, we applied linear mixed models (LMMs) to test traits we measured between *Col* and each mutant separately, considering genotypes as fixed effects and tray numbers as random effects. To account for false positives arising from multiple LMMs within the dataset, we calculated adjusted *p*-values using a False Discovery Rate (FDR) threshold of 0.05, applying the *p.adjust()* function in R. We summarized how genes, originally identified in previous studies, were selected for knockout experiments and examined whether they exhibited significant genotype effects. Additionally, we calculated the ratio of genes tested with significant genotype effects relative to the total number of genes selected for screening in each original study. We also applied LMMs on the daily g_sw_ and Fv/Fm data measured in the ID experiment to detect if the mutants differed from *Col* in ecophysiology on any of the 10 measuring days.

It was not feasible to control moisture in the freezing experiment as we did in the drought follow-up because soil lost water much more slowly than it did in higher temperatures, so we used *DunnettTest()* function in the R package *DescTools* (Signorell et al., 2023) to conduct Dunnett’s Test on traits measured in the freezing experiment, using *Col* as control and test for differences of the two mutants to control separately.

### Population genetic variation at *WRKY38*

Because we found some putative LoF for *WRKY38* were relatively common in the 1001 Genomes accessions, we further analysed the geographic and genetic structure of these variants. We used the imputed SNP matrix (hdf5 file) and short indels (vcf file) downloaded from the 1001 Genome Project website (https://1001genomes.org/accessions.html) for the following analyses.

We calculated Tajima’s D across chromosome 5 (Chr 5) in 5-kb windows using VCFtools (--TajimaD 5000) to test possible selection signals on *WRKY38* in natural accessions. The most common functional variant for *WRKY38* is a frameshift at position 7495793 on Chr 5 (Frameshift_7495793_; Table S3). To detect linkage disequilibrium and potential haplotype structure (i.e. non-random associations among SNPs within the genomic region) between Frameshift_7495793_ and adjacent SNPs, we calculated the square of the Spearman’s correlation coefficient (*r^2^*). This calculation was performed between the presence/absence of Frameshift_7495793_ and the alleles of each SNP within 5 kb upstream and downstream of *WRKY38* using the *cor()* function in base R.

To characterize the geographic and population structure of natural putative LoF mutations in 1001 Genomes accessions, we plotted mutations and their genetic clusters by different shapes and outline colors on a map. We also constructed neighbor-joining (NJ) trees built with the entire gene sequence of *WRKY38* (including coding and non-coding sequences), and a genome-wide NJ tree. We referred to the population structure of 1001 Genomes accessions in Alonso-Blanco et al. (2016), where the authors classified 9 genetic clusters and one admixed group. For simplicity of data presentation, we combined the 9 groups by 5 major regions — West Mediterranean (W-Med), North Mediterranean (N-Med), Central and Western Europe (CW-Euro), North Europe (N-Euro), and Asia. Accessions that were introduced, classified in the admixed group, or might be potential contaminations (Pisupati et al., 2017) were not included in the 5 groups.

We used the R package *maps* to visualize the distribution of the common putative LoF mutations of *WRKY38*. To build the genome-wide tree, we annotated the vcf file using SnpEff (Cingolani et al., 2012), extracted the synonymous sites, filtered out sites with minor allele frequency below 0.01 (--maf 0.01) and missing rate over 10% (--max-missing 0.9), and generated a distance matrix using PLINK (Purcell et al., 2007). Both NJ trees were constructed using the *nj()* function from package *ape* (Paradis and Schliep, 2019).

We hypothesized that the natural *WRKY38* putative LoF variants could underlie local adaptation. To identify which climate variables might best explain *WRKY38* functional variation and be most closely tied to mechanisms of selection, we used LMMs to scan for correlations across the 19 bioclimatic variables. Worldclim 2 bioclimatic data were retrieved using the *getData()* function from the R package raster (Fick and Hijmans, 2017; Hijmans et al., 2015). Models were implemented using *coxme* and *kinship2* packages to include kinship in the models as random effects (Sinnwell et al., 2014; Therneau and Therneau, 2015). We focused on testing the local adaptation of intact *WRKY38*, Frameshift_7495793_, the most common and widespread FV, and Frameshift_7495793&94_, which accounted for a 3-bp frameshift that led back to the normal open reading frame (ORF) thus suspected to be potentially functional. Since mixed models have low power when causal variants are correlated with the genomic background, we also used Welch’s t-test (*t.test()*) in R to compare the differences in bioclimatic variables for accessions with/out the intact gene or the FVs.

### Variation in *LSD1* expression

Only 5 of the total 1135 accessions had *LSD1* putative LoF mutations. Thus, we suspected that LoF may not be the major cause of the strong association between climate and the *LSD1*-targeting SNP in Lasky et al. (2018), and the adaptation might exist due to amino acid changes or *cis*-regulatory variation impacting expression. Therefore, we scanned across 19 bioclimatic variables and SNPs (ref/alt alleles) within 10kb around *LSD1* using univariate GEAs with GEMMA (Zoubarev et al., 2012) for 933 native accessions with known coordinates that were not potential contaminants (Pisupati et al., 2017). We found elevated correlations with climate at SNPs 1kb upstream of *LSD1* and the start of the gene sequence, suggesting that the adaptation may occur at the promoter region. These SNPs may influence gene expression by altering transcription factor binding motifs (Shastry, 2009; Wang et al., 2005).

Thus, we further asked if these SNPs impact *LSD1* expression using the published 1001 Genomes transcriptome data (http://signal.salk.edu/1001.php; Kawakatsu et al., 2016). Among 728 accessions with published transcriptome data, 1 accession was excluded from the following analysis according to Pisupati et al. (2017). We compared the expression levels of *LSD1* between the two alleles of the top climate-associated SNPs in the putative promoter region by Welch’s t-test (*t.test()*) in R. We also used linear models (*lm()*) to find possible associations between *LSD1* expression and bioclimatic variables. Additionally, we investigated whether SNPs with elevated correlations to climate are enriched in bioclimatic variables associated with *LSD1* expression.

## RESULTS

### Widespread putative functional variation in the experimentally screened genes in the 1001 Genomes accessions

Across all 1135 accessions, 1070 exhibit at least one high-impact functional variant (i.e. putative LoF) over the 44 experimentally screened genes. Specifically, 1973 putative frameshifts occurred in 943 genotypes as well as 1116 stop-gained FVs in 842 genotypes, appearing to be the most prevalent types of putative high-impact variants over the identified genes (Table S3). Splice donor variants were found in 303 accessions, and splice-acceptor, stop-lost, and start-lost variants were found in 29, 5, and 4 accessions, respectively.

Putative frameshifts were most common in AT1G19410 (*FBD*), occurring in 639 accessions. Frameshift variants were also common in AT5G22570 (*WRKY38*), detected in 250 accessions. Stop-gained had the most counts in AT5G22900 (*CHX3*), with 492 accessions having the putative FVs. Splice-donor was also the most prevalent in the *FBD* gene, exhibited by 220 accessions. The rest of the high-impact effects were rare among the screened loci, suggesting purifying selection on amino acid sequences. We found 14 accessions with splice-acceptor variants, 2 accessions with stop-lost, and 2 accessions with start-lost at AT5G10960 (*CAF1l*), AT3G45860 (*CRK4*), and AT5G22570 (*WRKY38*), respectively.

Some mutations in the screened genes had potential dual effects, and such dual effects were locusspecific. A 1-bp deletion or insertion at the same position in AT1G76090 (*SMT3*) caused both frameshift and start-loss, resulting in the most common dual effects observed in 95 accessions. Two insertions in AT3G45860 (*CRK4*) could each lead to dual effects of frameshift and stop-gain, together occurring in 21 accessions. In rare cases, a 1-bp deletion in AT5G40830 (*ICA*) caused frameshift and stop-lost in 4 accessions, while a 3-bp deletion in AT5G65050 (*AGL31*/*MAF2*) caused stop-lost and disruptive-inframe-deletion effects in 1 accession.

### Prevalent treatment effects, variable genotype effects, and few GxE effects on fitness and phenology in the main drought screen

Drought treatment effects on phenotypes were strong and largely consistent across different t-DNA insertion lines (Figure 1). Specifically, the drought effects were significant with FDR=0.05 across all 44 tested lines, causing lower aboveground biomass, smaller inflorescence length and weight, fewer siliques, shorter average and total silique length, and delayed flowering, with line CS68738 (*lsd1*) being the only exception with no significant drought effect on flowering time (Figure 1). The drought effect on rosette weight was also common, significantly reducing rosette weights in 25 of 44 lines at FDR = 0.05.

**Figure 1.**
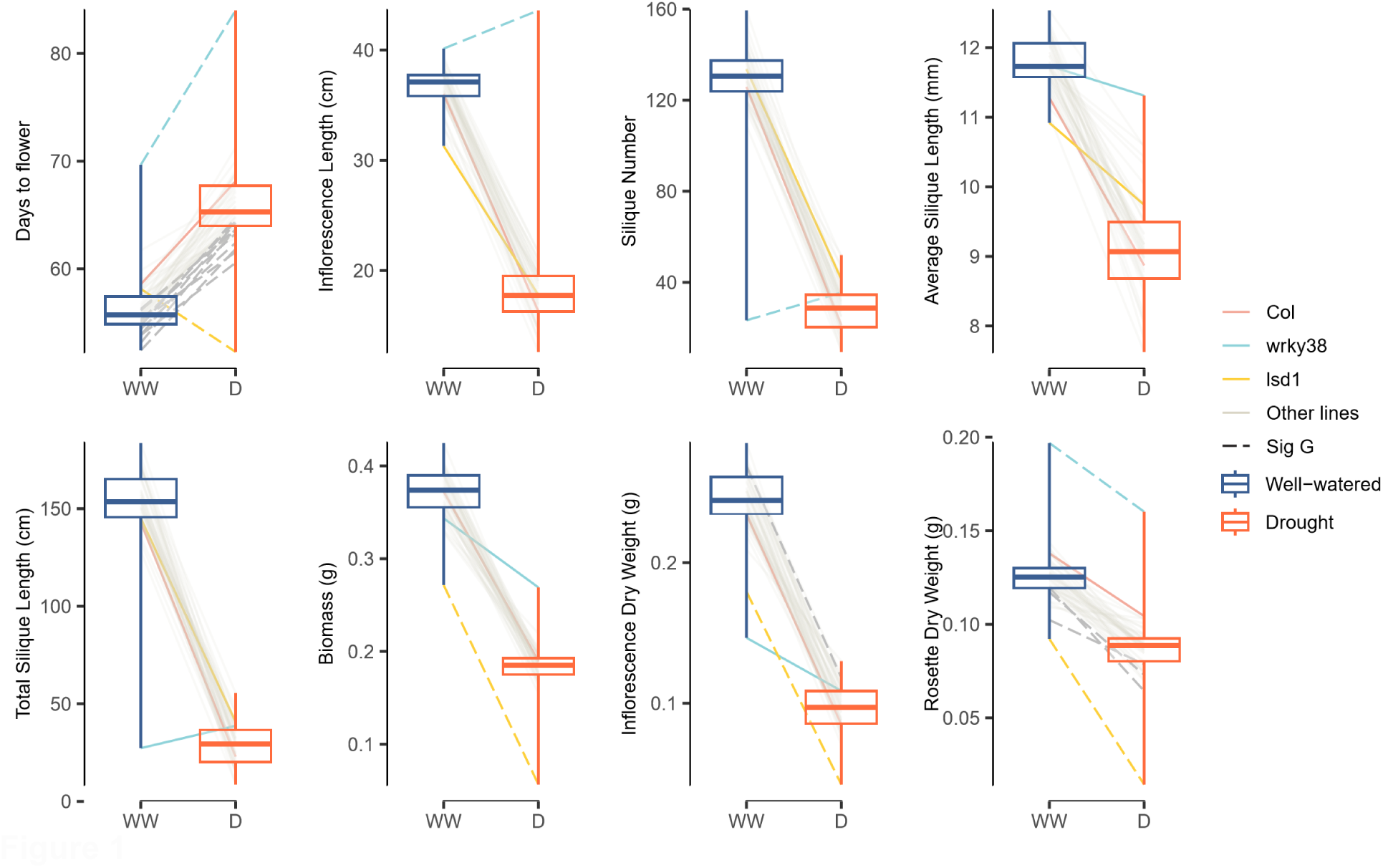
Reaction norms (lines) and average trait values (boxes) of a subset of traits measured during the main drought screen across all 44 mutants and *Col*, with *Col*, *lsd1* (CS68738), and *wrky38* (CS864818) highlighted. Each line connects the average trait values of one genotype under well-watered (WW) and drought (D) treatments, with *Col*, *lsd1* (CS68738), and *wrky38* (CS864818) highlighted. Mutants showing significant genotype effects at FDR = 0.05 from linear mixed models (LMMs) comparing *Col* and one mutant are indicated by dashed lines.

Genotype effects varied across the 44 lines tested (Figure 1, Table S4-6). Genotype effects were most common for flowering time, with 20 having nominally significant effects (α = 0.05) on flowering, primarily accelerating flowering, except for *wrky38*, which exhibited delayed flowering under drought. Thirteen insertion lines were significant with FDR = 0.05 (12 accelerated flowering and 1 delayed flowering (*wrky38*). Rosette and inflorescence weights were the traits with the next most insertion lines with genotype effects, 14 lines having nominally significant effects and 5 significant at FDR = 0.05 that mostly led to increased rosette weight and reduced inflorescence weight. Genotype effects on other traits measured were less common, with 5,5,5,3,1 lines having nominally significant effects and 1,0,0,1,1 lines significant with FDR = 0.05 for silique number, average and total silique length, inflorescence length, and aboveground biomass, respectively.

We found few genotype-by-treatment interactions caused by the t-DNA insertions. CS864818, a *WRKY38* (AT5G22570) knockout line, was the only one having a GxE effect that was significant at FDR = 0.05 on inflorescence length, silique numbers, total silique length, aboveground biomass, and inflorescence weight (Table 1,S5-S6). Generally, *wrky38* showed less drought sensitivity than *Col* while being smaller and less fecund in well-watered conditions except that the mutant had slightly longer inflorescence lengths than *Col* (*wrky38*: 40.1±3.010 cm; *Col*: 36.0±0.713 cm; Figure 2). When considering only genotype effects, the knockout of *WRKY38* caused significantly later flowering, longer inflorescence, more siliques, and greater rosette weights (Figure 2, Table 1,S4-S6. However, under drought conditions, only flowering time and inflorescence length were significantly different from *Col*.

**Figure 2.**
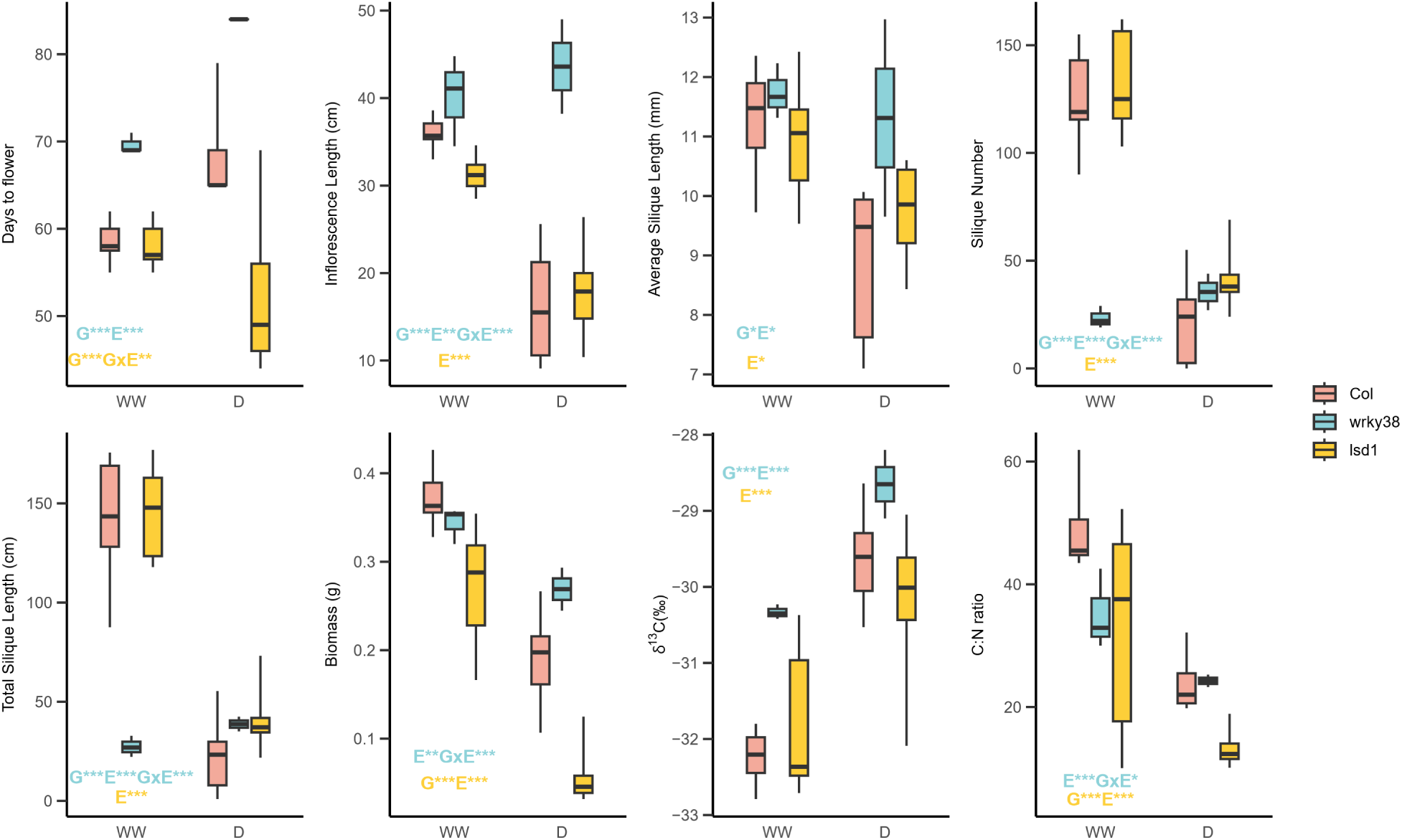
A subsetof the traits measured during the main drought screen of *Col*, *lsd1* (CS68738), and *wrky38* (CS864818). Text annotations on the box plots indicate the significance of genotype (G), treatment (E), and genotype-treatment interaction (G×E) effects at α = 0.05, based on LMMs comparing *Col* and one mutant. The text color denotes the mutant genotype. Significance levels are represented as follows: *, 0.01≤p<0.05; **, 0.001≤p<0.01; ***, p<0.001.

**Table 1.** Summary statistics from mixed linear models (LMMs) assessing a subset of performance and fitness traits for *Col*, CS864818 (*wkry38*) and CS68738 (*lsd1*) under well-watered and drought conditions during the main drought screen. Each LMM independently analyzes *Col* and one mutant (either *wrky38* or *lsd1*). β represents the estimated effect size, and subscripts indicate effects attributed to genotype (G), treatment (Trt) or genotype-by-treatment interaction (GxTrt); *p* represents the *p*-value at α = 0.05, and *p*_adj_ represents the adjusted *p*-value at FDR = 0.05. A negative β_G_ reflects a lower trait value in the mutant compared to *Col*, while a positive β_Trt_ reflects a lower trait value under drought compared to well-watered conditions.

| *Col* -CS864818 (*wkry38*) | Genotype Effect |  |  | Treatment Effect |  |  | Genotype-by-Treatment |  |  |
| --- | --- | --- | --- | --- | --- | --- | --- | --- | --- |
| | $\beta_G$ | *p* | *p*<sub>adj</sub> | $\beta_{Trt}$ | *p* | *p*<sub>adj</sub> | $\beta_{G \times Trt}$ | *p* | *p*<sub>adj</sub> |
| Days to Flower (days) | 15.86 | **<0.0001** | **0.0002** | -9.57 | **<0.0001** | **<0.0001** | -4.76 | 0.2380 | 0.8081 |
| Inflorescence Length (cm) | 27.33 | **<0.0001** | **0.0014** | 19.74 | **0.0086** | **0.0086** | -23.21 | **0.0006** | **0.0274** |
| Total Silique Length (cm) | 15.82 | **0.0018** | 0.0782 | 120.03 | **0.0009** | **0.0009** | -131.47 | **0.0002** | **0.0076** |
| Biomass (g) | 0.08 | 0.6793 | 0.9218 | 0.18 | **0.0059** | **0.0059** | -0.14 | **0.0004** | **0.0189** |
| *Col* -CS68738 (*Isd1*) | Genotype Effect |  |  | Treatment Effect |  |  | Genotype-by-Treatment |  |  |
| | $\beta_G$ | *p* | *p*<sub>adj</sub> | $\beta_{Trt}$ | *p* | *p*<sub>adj</sub> | $\beta_{G \times Trt}$ | *p* | *p*<sub>adj</sub> |
| Days to Flower (days) | -15.86 | **0.0009** | **0.0082** | -9.57 | 0.3981 | 0.3981 | 15.43 | **0.0015** | 0.0675 |
| Inflorescence Length (cm) | 19.74 | 0.3586 | 0.6422 | -6.20 | **<0.0001** | **<0.0001** | 20.14 | 0.0852 | 0.9698 |
| Total Silique Length (cm) | 17.75 | 0.2947 | 0.6362 | 120.03 | **<0.0001** | **<0.0001** | -15.68 | 0.4049 | 0.9684 |
| Biomass (g) | -0.13 | **<0.0001** | **<0.0001** | 0.18 | **<0.0001** | **<0.0001** | 0.03 | 0.4058 | 0.7440 |

Note: *p*-values less than 0.05 are presented in bold to indicate statistical significance.

Another mutant, *lsd1*, was one of only two having nominally significant genotype-by-treatment interaction (GxTrt) on flowering time (p = 0.0015; the other line SALK_012432 had p = 0.0479). The mutation significantly accelerated flowering compared to *Col*, and its interaction with drought exacerbated early flowering (Table 1). Notably, the mutant flowered earlier than *Col* under both WW and drought conditions (WW: 58.1±1.06 vs. 58.6±0.972 days; drought: 52.3±3.58 vs. 68.1±1.94 days), and unlike *Col* the effect of drought on flowering time was not significant (p = 0.3981). Drought had significant effects on fitness-related traits that reduced silique numbers, average silique length, and total silique length while the GxTrt effects on fitness were generally not significant (Table 1,S4-S6). lsd1 plants had significantly lower inflorescence and rosette weights, thus lower aboveground biomass, compared to *Col* (Figure 2, Table 1,S4-S6).

Both genotypes had increased WUE under drought compared to WW conditions (Figure 2), with treatment effects on WUE significant in LMMs that include *Col* and each mutant (p*_wrky38_* = 9.33e-18, p*_lsd1_* = 1.15e-6). The genotype effect of *WRKY38* knockout also significantly increased its WUE compared to *Col* (p = 8.81e-8), while that of *lsd1* was not significant (p = 0.756; Figure 2). *wrky38* had an increased δ^15^N ratio under drought compared to the well-watered condition, but its responses under both conditions were not substantially different from *Col* (p_G_ = 0.5010, p_Trt_ = 0.5575; Figure 2). *lsd1* mutants, however, had δ^15^N being lower under drought while slightly higher under WW compared to *Col* (p_G_ = 0.0387, p_GxTrt_ = 4.22e-4), suggesting a possible nitrogen source shift under drought. C:N ratios had been overall reduced under drought, probably the result of reduced biomass under drought (Figure 2).

### Comparing candidates from different GEA approaches

We compared each approach in identifying genes that influence fitness and performance traits under drought, potentially indicating their role in drought adaptation. Among the 40 top-candidate genes from the original studies, 12, 12, 8, 6, and 7 genes were identified through RDA, partial RDA, univariate associations, the combined common garden approach with aridity index, and the combined common garden approach with monthly growing season precipitation, respectively (Table S1). Four genes were identified as top candidates by multiple approaches (Table S1). Among the knockout lines of these genes, only SALK_051689 (AT5G10960) had marginally significant genotype effect that accelerated flowering time (p_adj_ = 0.045, Table S5).

RDA and partial RDA proved to be the most effective methods for identifying drought-adapted genes, with each approach having 6 out of 12 genes exhibiting significant genotype effects at FDR = 0.05 across fitness and performance traits in the main drought screen. The combined common garden approach followed, with 2 out of 6 and 2 out of 7 genes identified via association with aridity index and monthly precipitation showing significant genotype effects at FDR = 0.05.

In contrast, none of the 8 genes identified by univariate associations showed significant genotype effects at FDR = 0.05, although 5 genes exhibited significant genotype effects at α = 0.05 before applying false discovery rate corrections.

### Stable but less pronounced fitness, phenology, and ecophysiological responses of *wrky38* and *lsd1* under follow-up mild drought

Under the milder follow-up drought (ID), *wrky38* and *lsd1* plants also exhibited similar but less pronounced responses compared to those observed in the main drought screen. The mild drought significantly reduced fitness and inflorescence length across both *Col* and the mutants, as determined by mixed linear models that included *Col* and one mutant (*wrky38* or *lsd1*).

Specifically, the drought significantly reduced fitness and inflorescence length for both *wrky38* and *lsd1*, while flowering time was not significantly impacted (Figure 3, Table 2,S7). Plants of *wrky38* mutants did not exhibit significant genotype-specific responses to drought compared to *Col* across the traits we measured (Table 2). However, the direction of *wrky38* differences was consistent with the earlier drought screen; e.g., *wrky38* had higher fitness and biomass than*Col* under drought, but the opposite under well-watered conditions (Figure 3). In contrast, *lsd1* mutants showed significant genotype effects that reduce inflorescence length and total silique length and a significant GxE effect for silique number (Table 2). This indicates that *lsd1* mutants not only have reduced fitness under drought conditions, but the negative impact of drought was even more pronounced in *lsd1* mutants compared to *Col*.

**Figure 3.**
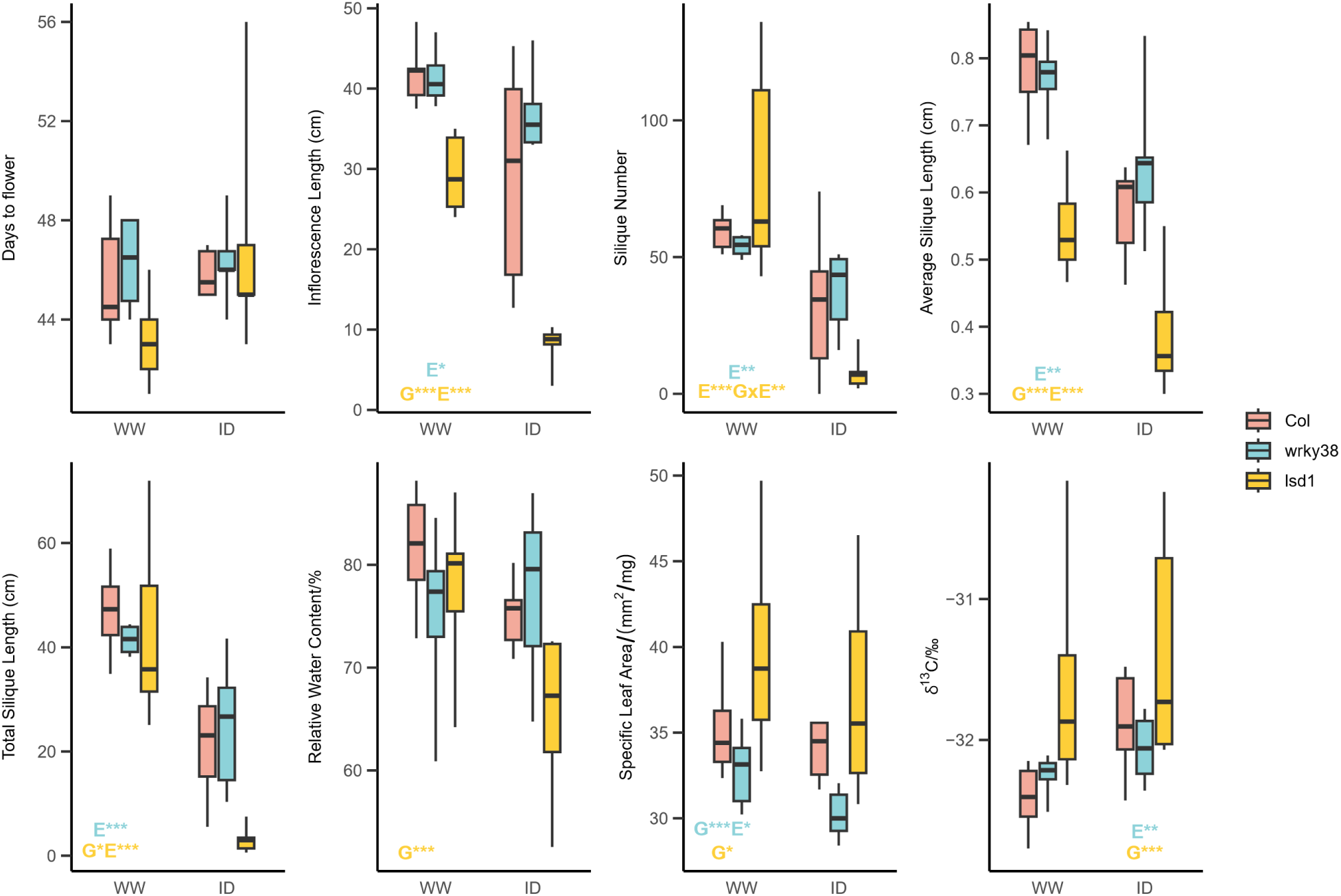
Traits measured in the follow-up intermediate drought experiment of *Col*, *lsd1* (CS68738), and *wrky38* (CS864818). Text annotations on the box plots indicate the significance of genotype (G), treatment (E), and genotype-treatment interaction (G×E) effects at α = 0.05, based on LMMs comparing *Col* and one mutant genotype. The text color denotes the mutant genotype. Significance levels are represented as follows: *, 0.01≤p<0.05; **, 0.001≤p<0.01; ***, p<0.001.

**Table 2.** Summary statistics from mixed linear models (LMMs) assessing a subset of performance and fitness traits for *Col*, CS864818 (*wkry38*) and CS68738 (*lsd1*) under well-watered and intermediate drought (ID) conditions during the follow-up intermediate drought experiment. Each LMM independently analyzes *Col* and one mutant (either *wrky38* or *lsd1*). β represents the estimated effect size, *p* represents the *p*-value at α = 0.05, and subscripts indicate effects attributed to genotype (G), treatment (Trt) or genotype-by-treatment interaction (GxTrt). A negative β_G_ reflects a lower trait value in the mutant compared to *Col*, while a positive β_Trt_ reflects a lower trait value under ID compared to well-watered conditions.

| *Col* -CS864818 (*wkry38*) | $\beta_G$ | *p*<sub>G</sub> | $\beta_{Trt}$ | *p*<sub>Trt</sub> | $\beta_{G \times Trt}$ | *p*<sub>G × Trt</sub> |
| --- | --- | --- | --- | --- | --- | --- |
| Days to Flower (days) | 0.50 | 0.1503 | -0.33 | 0.9684 | 0.93 | 0.4626 |
| Inflorescence Length (cm) | 7.75 | 0.2416 | 12.67 | **0.0105** | -8.11 | 0.2551 |
| Silique Number | 5.00 | 0.9632 | 26.67 | **0.0023** | -10.50 | 0.4705 |
| Total Silique Length (cm) | 3.63 | 0.8752 | 25.68 | **<0.0001** | -9.23 | 0.2925 |
| Specific Leaf Area (mm<sup>2</sup>/mg) | -3.80 | **0.0005** | 1.19 | **0.0313** | 1.43 | 0.4181 |
| *Col* -CS68738 (*Isd1*) | $\beta_G$ | *p*<sub>G</sub> | $\beta_{Trt}$ | *p*<sub>Trt</sub> | $\beta_{G \times Trt}$ | *p*<sub>G × Trt</sub> |
| Days to Flower (days) | 1.42 | 0.7466 | -0.33 | 0.1908 | -3.63 | 0.1365 |
| Silique Number | -25.00 | 0.8293 | 26.67 | **0.0007** | 48.51 | **0.0081** |
| Inflorescence Length (cm) | -21.10 | **<0.0001** | 12.67 | **<0.0001** | 8.65 | 0.1897 |
| Total Silique Length (cm) | -18.25 | **0.0234** | 25.68 | **<0.0001** | 14.45 | 0.1378 |
| Specific Leaf Area (mm<sup>2</sup>/mg) | 3.11 | 0.0368 | 1.19 | 0.4479 | 1.41 | 0.6989 |

Note: *p*-values less than 0.05 are presented in bold to indicate statistical significance.

During the 10 days of ecophysiological tracking, we built LMMs to compare the daily *g_sw_* and Fv/Fm of the two mutants with *Col*. The stomatal conductance (*g_sw_*) of all three genotypes exhibited fluctuations that were associated with the regular watering schedule (Figure S1). *wrky38* had potentially adaptive reduced *g_sw_* compared to *Col* and under drought, with genotype effects significant during the initial days of tracking, when drought-treated plants first experienced water limitation (9 days after treatment initiation; Table S3). A significant GxE interaction with drought was also observed (Table S8). The *lsd1* mutant also had reduced *g_sw_* under drought, but the genotype effect was insignificant (Figure S1; Table S8). None of the Fv/Fm values we measured suggested major oxidative stress (Figure S2), coinciding with Zivcak et al. (2013), indicating a mild drought. No genotype or treatment pattern was detected for Fv/Fm.

The drought effect significantly reduced SLA when considering *Col* and *wrky38* but was not significant in the model including *Col* and *lsd1* (Figure 3, Table 2). Genotype effects on SLA were significant for both mutants, but *wrky38* had a lower SLA (potentially adaptive under drought) while *lsd1* had a higher SLA compared to *Col* (Figure 3, Table 2). RWC exhibited little variation between genotypes and treatments except for *lsd1* plants, which had a significant genotype effect resulting in lower RWC (Table S7). The δ^13^C response to treatment was generally lower in the follow-up than in the main drought screen, indicating a milder drought in the follow-up. The treatment effect on WUE was only significant for *wrky38* (p = 0.0029), whereas the genotype effect on WUE was only significant for *lsd1* (p = 3.7e-4). No significant GxE effects were detected for SLA, RWC, or δ^13^C (Table S7).

### Reduced vegetative growth or fitness of *wrky38* and *lsd1* under overnight freezing

Treated with overnight freezing for 30 days, both *lsd1* and *wrky38* had significantly smaller diameters than *Col* (p*_wrky38_* = 0.0184, p*_lsd1_* = 3.7e-4), representing their reduced vegetative growth under low temperatures (Figure 4). After returning to warmer temperatures, we found no variation in flowering time and silique number between *Col* and the two mutants. However, *lsd1* had a shorter inflorescence length (p = 0.0360), average silique length (p = 0.0023), and total silique length (p = 0.0218) than *Col*. Both *lsd1* and *wkry38* had smaller biomass than *Col* (p*_wrky38_* = 0.0026, p*_lsd1_* = 2.6e-4), which might also be due to reduced vegetative growth. Overall, *lsd1* plants had reduced growth and fitness compared to *Col* under freezing, consistent with findings of Huang et al. (2010) that *lsd1* is chilling sensitive. Although putative LoF variants in *WRKY38* were strongly associated with warm winter temperatures (Lasky et al. 2018), *WRKY38* knockouts did not exhibit fitness differences compared to *Col*, though their vegetative growth was inhibited by overnight freezing compared to *Col*.

**Figure 4.**
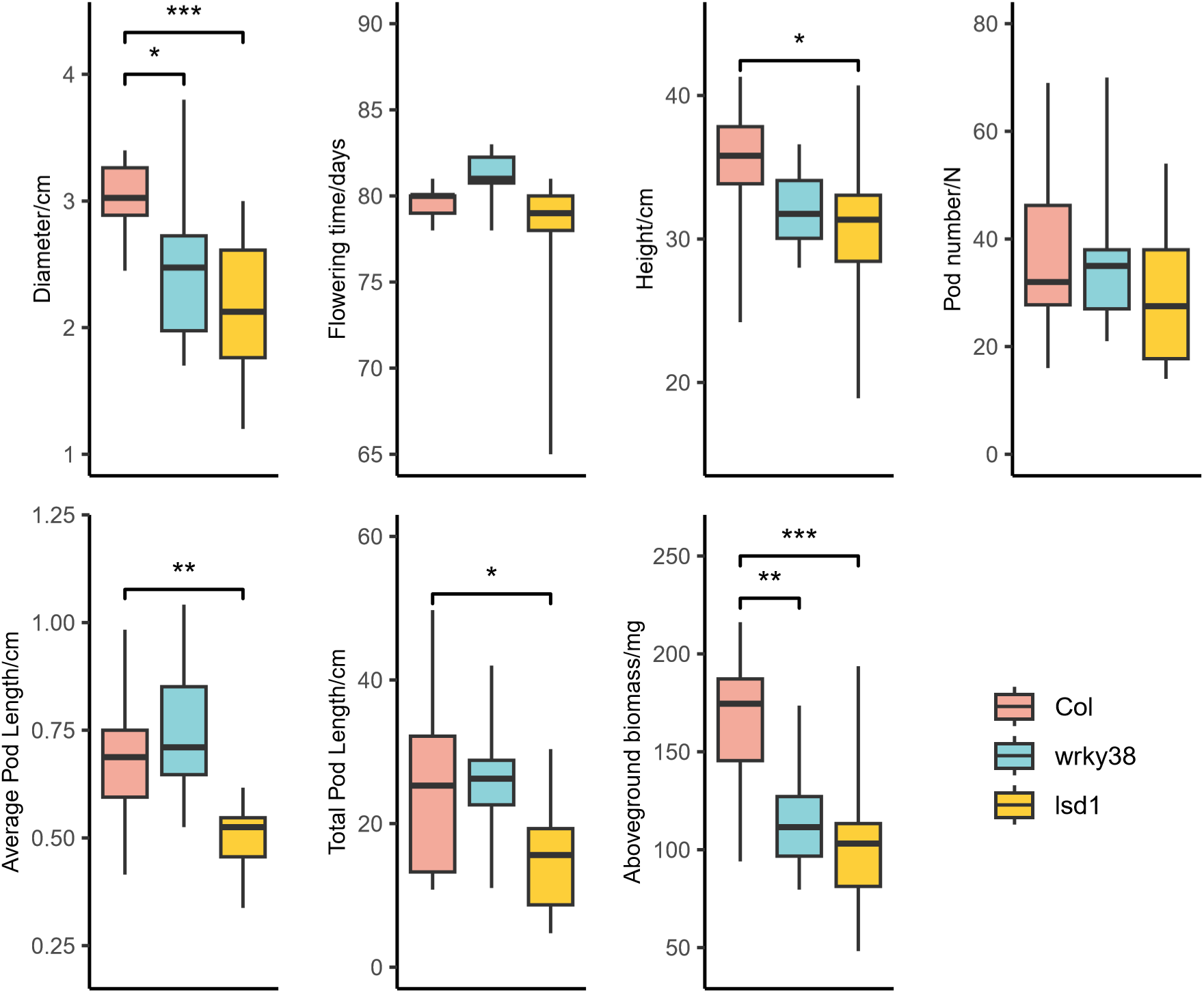
Traits measured in the follow-up freezing experiment of *Col*, *lsd1* (CS68738), and *wrky38* (CS864818). Asterisks indicate significant trait differences between *Col* and each mutant based on Dunnett’s test: *, 0.01≤p<0.05; **, 0.001≤p<0.01; ***, p<0.001.

### *WRKY38* allelic variation associated with climatic moisture and temperature

To evaluate potential selection on *WRKY38* and its surrounding genomic region, we analyzed SNP diversity patterns and examined allele associations indicating possible selection effects. The Tajima’s D of the 5-kb window containing the *WRKY38* gene was above 0 (Tajima’s D = 1.429) and is above the average of Chromosome 5 (average Tajima’s D = 0.660), but it did not significantly deviate from the distribution of Tajima’s D across the chromosome (*z-*score = 0.659; Figure S3). The overall *r^2^* between SNPs in 10 kb around *WRKY38* and the most common functional variant (Frameshift_7495793_) also indicated some haplotype structure, with the frameshift having elevated correlations (*r^2^* >0.4) for SNPs located within 2 kb upstream of *WRKY38* and in the first 500 bp of the gene (also where the frameshift was located, maximum *r^2^* = 0.5104; Figure S4). However, this was not a dramatically long region of linkage, suggesting no evidence for a recent sweep (cf. the strong correlation observed with chr4:10999188 and other SNPs in a 300 kb window around *LSD1,* Lee et al., 2017). Nevertheless, the putative LoF mutations carried large numbers of alternate SNP alleles across the locus, indicating some divergence and linkage (Figure 5i,S5). The haplotype structure at *WRKY38* can also be seen in the *WRKY38* gene neighbor-joining tree. While the genome-wide NJ tree shows a star-like structure consistent with the recent expansion of Arabidopsis (Lee et al. 2017) with accessions clustered based on geography (Figure 5f), the *WRKY38* tree shows strong divergence between sequences with Frameshift_7495793_ versus putative intact *WRKY38* sequences (Figure 5g,h). This allelic variation is segregated across diverse lineages, including the relicts (Figure 5a,b,c,f). Coincidentally, the rare FVs were all unique to their lineages, suggesting recent origins (Figure 5f; Table S9). In contrast to the widespread Frameshift_7495793_, they might reflect newer local putative LoF mutations.

**Figure 5.**
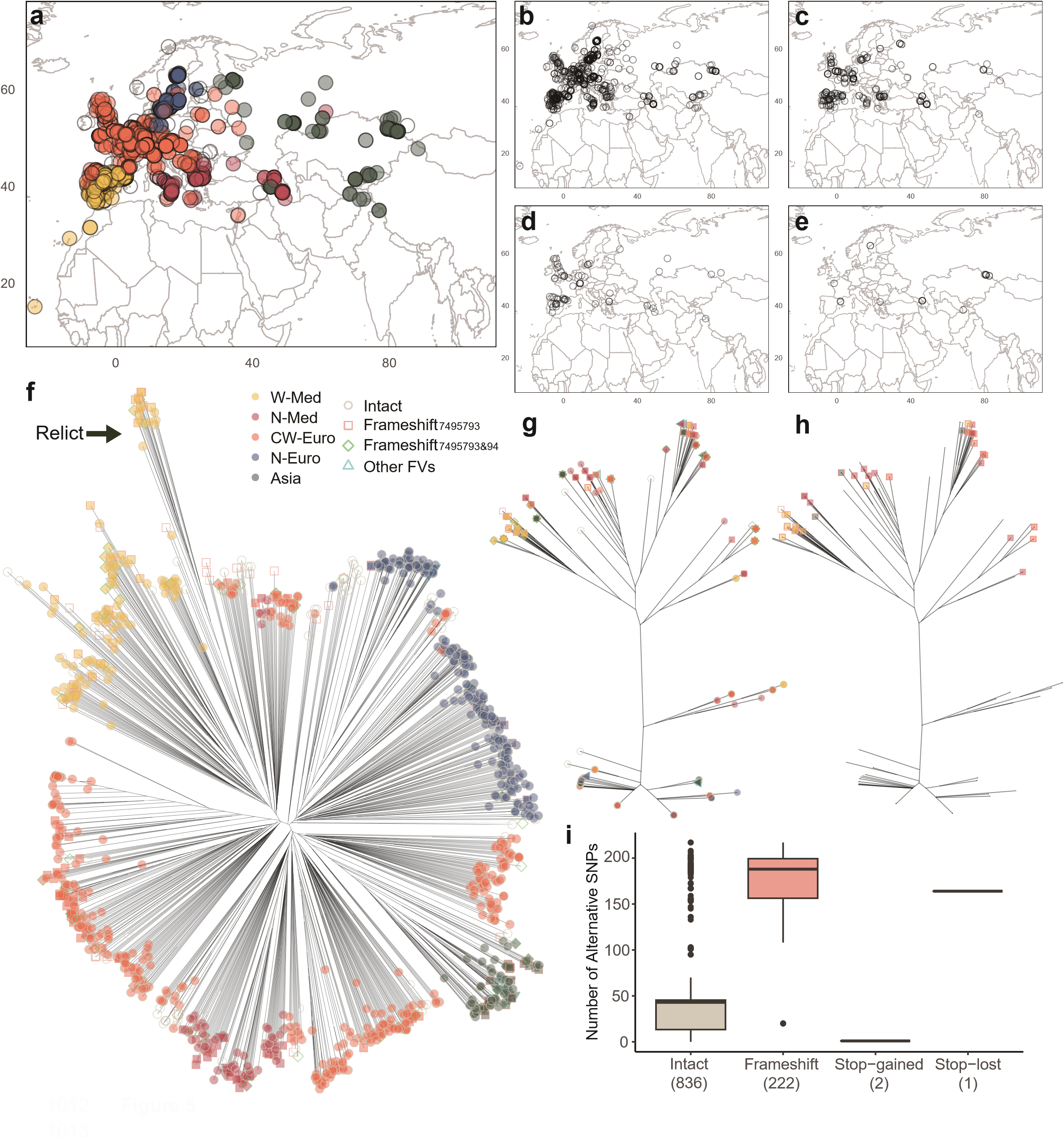
Natural variation of *WRKY38* putative LoF in the 1001 Genomes accessions. **a**, distribution of native *Arabidopsis* accessions from 1001 Genomes. Colors indicate ADMIXTURE genetic clusters, originally defined by Alonso-Blanco et al. (2016) and modified in this study. Accessions with clearly assigned genetic clusters are represented by filled icons in their respective cluster colors, while admixed accessions are shown with unfilled icons. **b**-**e**, distributions of natural *WRKY38* putative functional variation: **b**, distribution of accessions with intact *WRKY38*; **c**, distribution of accessions with Frameshift_7495793_, the most common FV of *WRKY38*; **d**, distribution of accessions with Frameshift_7495793&94_, a mutation that restores the normal open reading frame; **e**, distribution of accessions with other rare FVs of *WRKY38*. **f,** *WRKY38* FVs on the genomewide NJ tree, the relict lineage (Alonso-Blanco et al., 2016) was marked by an arrow. **g**, *WRKY38* FVs on the NJ gene tree, symbols as in (**f**). **h,** Frameshift_7495793_ on the NJ gene tree. **i**, numbers of alternative SNPs in accessions with intact *WRKY38* and putative LoF FVs within 10 kb around *WRKY38*. The number of accessions with each type of functional variation is indicated in parentheses.

The gene tree of WRKY38 shows that alleles with the same functional variants exhibit similar haplotype patterns, regardless of their genetic cluster (Figure 5g,h). For example, although most N-Euro accessions are in the non-Frameshift_7495793_ clade, all 12 N-Euro accessions having Frameshift_7495793_ are in the Frameshift_7495793_ clade, overlapping with W-Med, CW-Euro, and Asian accessions carrying this allele. The only accession having Frameshift_7495794_ and the only one with Stop-lost_7495609_ are both within Frameshift_7495793_ clade, while the 6 accessions having Frameshift_7496041_ and 2 accessions having Stop-gained_7496443_, which are grouped at the same tip on the gene tree, only occurred in the non-Frameshift_7495793_ clade.

Inconsistency between the genomewide and *WRKY38* gene tree confirmed *WRKY38* allelic variation independent of population genetic structure. Next, we investigated whether this variation is associated with climatic factors, particularly moisture and drought related variables. We did not find significant correlations at ⍺ = 0.05 between the frequency of Frameshift_7495793_ (the most common FV) or Frameshift_7495793&94_ (the FV that putative restores the normal ORF) and the bioclimatic variables when considering kinship (Table S10), which coincided with the findings of Alonso-Blanco et al. (2016) that almost no SNPs are significantly correlated with climate in environmental GWAS at FDR = 0.05 after considering population structure for 1001 Genomes accessions. Despite this, without accounting for kinship, significant differentiation was detected at ⍺ = 0.05 for 11, 12, and 4 bioclimatic variables between accessions with and without intact *WRKY38*, Frameshift_7495793_, and Frameshift_7495793&94_, respectively (Table S11). Among the significant associations, precipitation of the warmest quarter (bio18) had the lowest p-values for both intact *WRKY38* (p = 6.8e-10) and Frameshift_7495793_ (p = 8.8e-9). Specifically, intact *WRKY38* occurred more frequently in regions with wet summers, while Frameshift_7495793_ was more common in hot and dry regions, suggesting the drought adaptation of *WRKY38* putative LoF. Frameshift_7495793&94_ (which has a net in-frame effect) was most significantly correlated with higher isothermality (bio3; p = 3.7e-4), followed by higher mean temperature of the wettest quarter (bio8; p = 6.4e-4).

### *LSD1* allelic variation and expression associated with climatic moisture and temperature

Mixed models (GEMMA) detected strong correlations between SNPs within 10kb around *LSD1*, particularly for SNPs within 1 kb upstream and the first 500 bp of *LSD1*, and multiple bioclimatic variables (Figure 6a). These SNPs, located in the promoter region of *LSD1*, may influence gene expression by altering transcription factor binding motifs (Shastry, 2009; Wang et al., 2005). Thus, we hypothesized that these SNPs might contribute to local adaptation via changes in gene expression.

**Figure 6.**
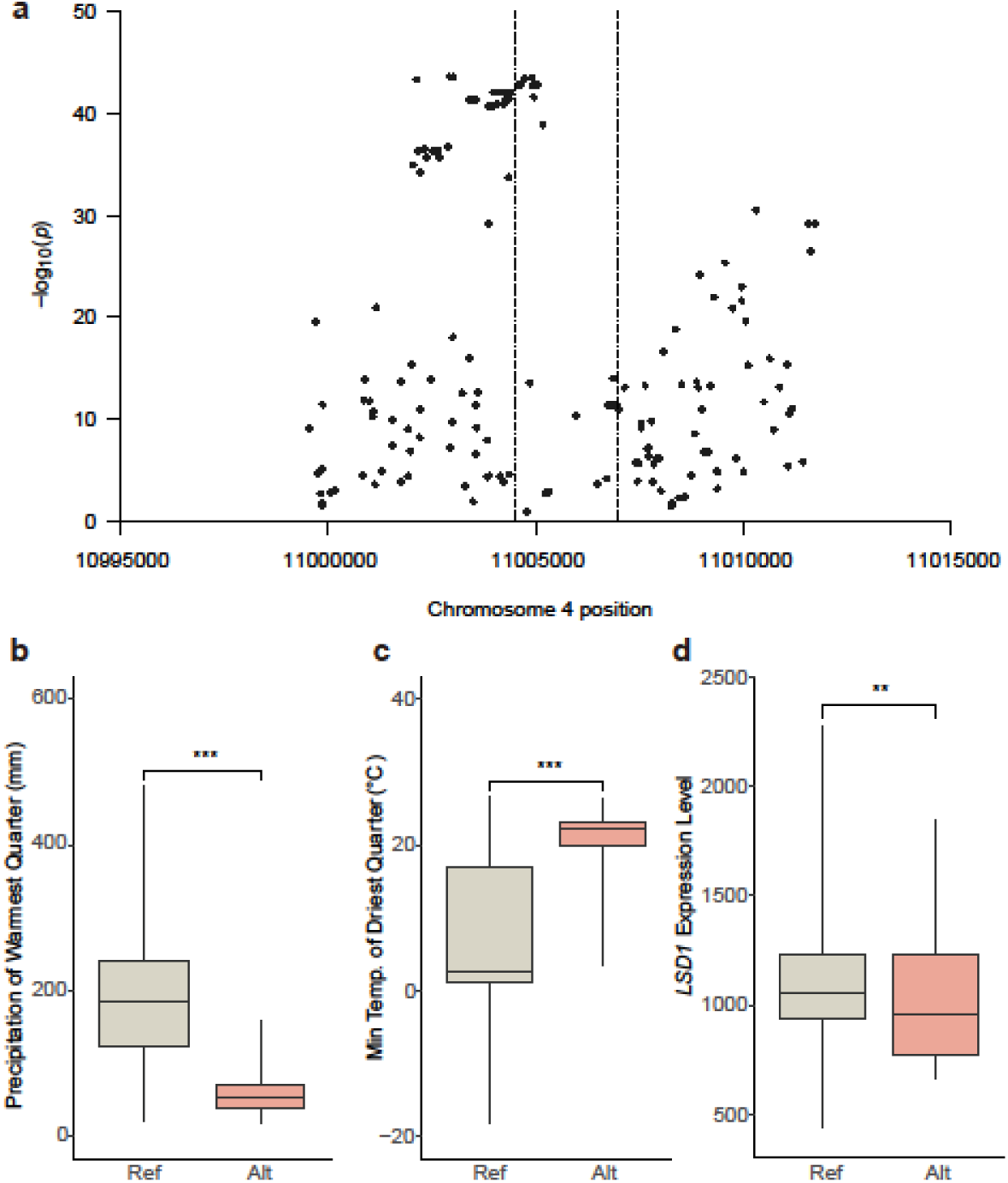
Genotype-environment associations and *LSD1* expression. **a**, Scanning the *LSD1* region to identify potential SNP-environment associations using LMM within 10kb. Each point represents the smallest p-value from the 19 associations between each SNP and the 19 bioclimate variables. *LSD1* gene region is shown between the dashed lines. These associations are meant to identify where in the locus variation is most strongly associated with climate. **b**, the difference in precipitation of the warmest quarter (bio18) between reference and alternative alleles at a top climate-associated SNP (Chr4, position 11003866). **c**, the difference in the minimum temperature of the driest quarter (bio9) at an SNP (Chr4, position 11003866). **d**, the difference in *LSD1* expression at an SNP (Chr4, position 11003866). **b-d**, asterisks indicate significant trait differences from t-test: *, 0.01≤p<0.05; **, 0.001≤p<0.05; ***, p<0.001.

To validate the hypothesis, we picked a top climate-associate SNP (Chr4, position 11003866) from the 1 kb upstream region of *LSD1* and tested for *LSD1* expression differences between ref/alt alleles. This SNP had the strongest correlation, indicated by the lowest p-value, with precipitation of the warmest quarter (bio18; wald-p = 2.390e-41; Figure 6b), followed by the minimum temperature of the driest quarter (bio9; wald-p = 2.140e-36; Figure 6c). We found that the alternative allele corresponded to significantly lower *LSD1* expression compared to the reference allele (p = 0.003; Figure 6d), suggesting a connection between lower *LSD1* expression and hot and dry climates.

Using linear models, we found that *LSD1* expression was significantly greater for genotypes from wetter summer climates (precipitation of the warmest quarter (p=8.683e-5) and precipitation of the wettest month (p=0.039)). Across the 10 kb region around *LSD1*, 68 of the 201 SNPs were most strongly associated with bio18 (precipitation of the warmest quarter).

Moreover, when limiting to SNPs within 1kb upstream of *LSD1*, i.e. the region we thought to be linked with *LSD1*, the ratio became 25 of 35 SNPs. The results indicated that *LSD1* might play a role in local adaptation to moisture regimes by *cis*-regulatory variation.

## DISCUSSION

Local adaptation is a major source of phenotypic variation within species, but traditional methods of identifying locally adapted genes are logistically challenging. Genotype-environment associations (GEA) offer an easy-to-implement tool to generate hypotheses about genes involved in local adaptation, but most published results remain as untested hypotheses. Here we screened 42 genes identified from genome-wide GEA studies and found at least two had effects on environmental responses and physiology linked to drought adaptation, suggesting potential local adaptations.

In our main drought experiment, among 42 potential drought-adapted genes detected with 3 different GEA approaches, the tDNA knockout of *WRKY38* (detected with RDA, Lasky et al., 2012) exhibited a significant GxE effect on fitness-related traits, and *lsd1* mutant (gene detected with combined GEA-common garden synthesis, Lasky et al., 2018) had a significant GxE effect on flowering time, which could potentially indicate variation in drought escape versus drought avoidance (Lovell et al., 2013). Coincidentally, these genes were both the top (#1) candidate genes with the strongest GEA in the studies used to select them. The physiological effect of *wrky38* identified in the follow-up screening suggests mechanisms of drought avoidance for *wrky38*, via reduced stomatal conductance and SLA. Furthermore, the high frequency of putative natural loss-of-function alleles of *WRKY38* in Arabidopsis populations found in the drier southern part of its range suggests potential local adaptation.

### Genotype and genotype-by-enviroment effects on phenotypes

Although the screened genes were the ones with the largest effects in the original studies, the GxE effects for fitness that were significant in the main drought screen were no longer significant under the milder drought conditions of the follow-up ID experiment. Several factors could impact the results. First, the milder conditions of the follow-up drought experiment compared to the main drought screen may have reduced the G×E effect on fitness. Second, over-simplified laboratory environments could lead to systematic bias in plant fitness (Anderson et al., 2013). In natural environments, drought conditions are usually coupled with elevated temperatures, a variable not accounted for in controlled experiments designed to isolate the effects of a single environmental factor. In the laboratory, when only a single environmental variable is changed between treatments, a conditional neutrality, that a gene has fitness advantages under one treatment and is neutral under the other, can be more common (Anderson et al., 2013). Lastly, the knockout alleles of a gene may not be the mechanism of local adaptation in nature. Local adaptation may also occur through *cis*-regulatory effects on gene expression (Hämälä et al., 2020; Lasky et al., 2014) or changes in amino acid sequences. Therefore, when screening candidate genes for local adaptation, it is important to consider lines with significant genotype effects alongside those showing significant G×E effects for fitness. As G×E effects may not always be significant due to the factors discussed above, genotype effects may still capture environment-associated differentiation.

Besides G×E effects on fitness, G×E effects on flowering time are also noteworthy when assessing local adaptation to drought stress, as early flowering is a critical drought escape strategy (Franks, 2011; Kenney et al., 2014). Chong and Stinchcombe (2019) reported systematic directional effects of flowering time among randomly selected SALK lines. In their study, 11 out of 27 lines under long-day conditions and 3 out of 27 lines under short-day conditions exhibited significantly different flowering times compared to *Col*, with all showing delayed flowering. However, in our main drought screen, 19 out of 20 lines with significant genotype effects at ⍺ = 0.05, and 12 out of 13 lines with significant genotype effects at FDR = 0.05, exhibited accelerated flowering (Figure 1, Table S6). Among these, *wrky38* was the only line that showed delayed flowering (a potential drought avoidance strategy, Ludlow, 1989). Since the genes we selected were candidates for drought adaptation, the accelerated flowering observed in these lines may indicate their potential for drought adaptation via escape (Ludlow, 1989).

Though we only observed GxE for fitness in a small number of genes, evidence of associations between their natural functional variation and moisture-related bioclimate variables demonstrated the underlying biological bases of potential local adaptation of loci identified by GEA. Setting up this screen was relatively easy due to the availability of existing tDNA resources, making the approach an efficient way to identify and screen potential locally adapted genes in model organisms where such resources are available or relatively easily generated.

### Comparison of common GEA approaches

Since the advent of GEA, a wide variety of methods have been proposed (reviewed by Rellstab et al., 2015). We tested candidates from three approaches here: multivariate ordination (RDA) of genotype and environment (Forester et al., 2018; Lasky et al., 2012), univariate climate mixed model associations (Lasky et al., 2014), and univariate climate models integrated with fitness data from multiple common gardens (Lasky et al., 2018). Even considering lines exhibiting significant genotype effects in addition to GxTrt effects, the number of positives remains too small to definitely compare these statistical GEA approaches. While Lotterhos (2023) recently criticized RDA as inappropriate for GEA due to a high false positive rate, the author used arbitrary and uncalibrated significance thresholds. By contrast, Figure S8 in Lotterhos (2023) showed that RDA was among the best methods based on the area under the precision-recall curve (AUC-PR). Our results further support this, showing that RDA and partial RDA had the highest rate of genes exhibiting significant genotype effects at FDR = 0.05, followed by the combined common garden approach. In comparison, none of the tested genes identified through univariate associations showed significant genotype effects at FDR = 0.05.

We propose that using ranked GEA results is more informative than focusing on arbitrary significance thresholds; as mentioned previously, *WRKY38* and *LSD1* were the top-ranked loci in their respective studies (RDA and combined GEA-common garden mixed models, respectively; Lasky et al. 2012, 2018).

### Potential mechanisms of drought adaptation at *WRKY38* and *LSD1*

The two mutants presented opposite responses under drought. *wrky38* had a significantly smaller SLA when compared to *Col* and when under drought, indicating how *wrky38* maintained stable RWC under well-watered and drought conditions. *wrky38* mutants have reduced g_sw_, SLA, and leaf area (result not shown) under drought in the follow-up experiment and increased WUE in response to drought during both experiments (Figure 2-3), suggesting a drought-avoidance strategy of natural *WRKY38* LoF genotypes. In contrast, *lsd1* had a larger SLA than *Col*, thus potentially more water loss (Niinemets, 2001), which may result in its reduced RWC. Instead, *lsd1* mutants showed stable accelerated flowering under drought in our two drought experiments, indicating a drought-escape strategy.

*WRKY38* negatively regulates plant basal pathogen defense by suppressing *PATHOGENESIS-RELATED GENE 1* (*PR1*) expression, a defense gene induced by salicylic acid (SA) (Kim et al., 2008). Additionally, *WRKY38* is also involved in drought- and cold-related responses in barley (Marè et al., 2004). Among *Arabidopsis WRKY* members that were also acting in the SA pathway for plant basal defense responses (Salinas et al., 2024), *AtWRKY54* and *AtWRKY70* were found to negatively impact osmotic tolerance by reducing stomatal closure (Li et al., 2013), while *AtWRKY46* was found to regulate light-dependent stomatal opening in guard cells (Ding et al., 2014). Based on previous studies and our findings, we hypothesize that *WRKY38* inhibits stomatal closure. If so, the lower *g_sw_* we found in knockouts could reduce water loss through the stomata and enhance fitness under drought.

In 1001 Genome accessions, natural *WRKY38* LoF alleles were strongly associated with lower July moisture compared to intact alleles. Despite this strong association, these putative LoF alleles were not fixed or entirely absent in any region. Our population genetics analyses revealed that LoF *WRKY38* exists across the entire distribution region of *A. thaliana* and in every lineage group (Figure 5a-f). We suspected that the LoF of *WRKY38* might have originated before the migration across Eurasia of *A. thaliana* and colonized drier regions during this process. In both drought experiments, *wrky38* generally had lower fitness compared to *Col* under well-watered conditions. However, *wrky38* maintained relatively stable fitness under drought stress, while *Col* showed a substantial fitness decrease. The drought insensitivity of *wrky38* may explain how natural LoF *WRKY38* dominates drier regions during migration.

The absence of LSD1 protein in *lsd1* mutants promotes the accumulation of SA (Bernacki et al., 2021), which adversely affects plant development and reproduction while accelerating flowering in Arabidopsis (Salinas et al., 2024). These findings explained the consistent accelerated flowering behavior observed across our two drought experiments. However, *lsd1* plants differed in their fitness responses to the long-term water deficiency of our main drought screen and periodical moisture fluctuation in our follow-up experiments (Figure 2-3). Specifically, *lsd1* mutants had similar fitness to *Col* under well-watered and drought in the main experiment, but they were less fit than *Col* (i.e. genotype effect was significant) in the follow-up. Previous laboratory studies have shown that *lsd1* exhibits similar survival and seed production to the wild type under nonlethal water deficiency. However, under lethal drought stress that caused complete mortality in wild-type plants, *lsd1* demonstrated a significantly higher survival rate, albeit with reduced seed production. These findings suggest a trade-off between survival and fecundity under severe stress in controlled laboratory conditions (Szechyńska-Hebda et al., 2016; Wituszynska et al., 2013). When grown in the field, *lsd1* mutants did not exhibit seed yield differences compared to the wild type (Bernacki et al., 2019; Szechyńska-Hebda et al., 2016; Wituszynska et al., 2013). This could explain our finding that the *LSD1* gene had lower expression in the hotter and drier regions, as reduced expression could similarly mitigate fitness trade-offs, enhancing survival in stressful climates.

### Freezing responses of *wrky38* and *lsd1*

We conducted the freezing experiment because both genes may also be involved in adaptation to freezing stress. A SNP within the *WRKY38* coding region (Chr. 5, pos:7496047) was the top candidate loci associated with temperature of the coldest month in Lasky et al. (2018), and *LSD1* is associated with chilling sensitivity (Huang et al., 2010). Moreover, these two genes are both in the salicylic acid (SA) pathway that can be activated by cold stress (Miura and Tada, 2014; Wu et al., 2019). *WRKY38* was positively regulated by SA (Kim et al., 2008) while *LSD1* conditionally regulated SA concentration with the existences of *ENHANCED DISEASE SUSCEPTIBILITY1* (*EDS1*) and *PHYTOALEXIN DEFICIENT4* (*PAD4*) (Bernacki et al., 2019; Szechyńska-Hebda et al., 2016; Wituszynska et al., 2013).

*WRKY38* knockouts exhibited similar fitness to *Col* despite reduced vegetative growth after long-term overnight freezing (Figure 4). Considering that *WRKY38* was originally identified through RDA (Lasky et al., 2012), this finding suggests a potential role in multivariate adaptation. The LoF allele might be advantageous in drier and warmer climates, while the functional allele might be more beneficial in cooler and wetter climates. *LSD1* knockouts had significantly reduced vegetative growth and fitness under freezing, coincident with the findings of Huang et al. (2010). Both mutants exhibited some disadvantages under freezing conditions compared to *Col*, indicating possible links between the function of these genes and adaptation to cold climates. Although both drought and freezing conditions can limit water availability and there is overlap in some molecular pathways or responses to each set of conditions (Kim et al., 2024), here our results suggest the drought-adaptive variants are not cold-adapted.

## Conclusion

In this study, we tested the potential role of 44 t-DNA knockout mutants of GEA-identified genes in adaptation to drought stress using common garden experiments. While most mutants did not exhibit significant G×E effects for flowering time, performance, or fitness, two knockouts, *wrky38* and *lsd1*, demonstrated evidence of drought adaptation in both the main drought screen and follow-up intermediate drought common gardens. Natural variation in the function or expression of these genes across a moisture gradient further supports the utility of GEA approaches for generating hypotheses, though further experiments are required to test hypotheses. Our findings highlight the promise of GEA methods for uncovering novel local adaptations to environmental stressors.

## Acknowledgments

We thank Cody Depew for assistance in ordering knockout lines, Amanda Penn for assistance with experiments, and Diana Gamba and Yuxing Xu for assistance with computation. Funding was provided by NIH R35GM138300 to JRL.

## Author contributions

YL, CML, and JRL designed the study, CL and YL performed experiments, YL conducted data analyses, YL, ELP, and JRL interpreted results, YL, CML, and JRL wrote the manuscript, all authors edited and approved the manuscript.

