## Supplementary material for "Experimental validation of genome-environment associations in Arabidopsis": Supp-Info-Table-of-Content.docx

**Table of Contents in Supplemental Information.zip:**

| **Supplementary Methods** | Page 1-4 |
| --- | --- |
| **Figure S1-S4 and captions** | Page 5-8 |
| **Figure S5 caption only** | Page 8 |
| **Table S1-S11 and captions** | See files of their name |

**Note:**

- Separate pdf files for each supplementary figure are provided in the zip file. Figure 5 is not included in the current file because of figure size.
- Figure captions are all provided in the current document.
- Table S1-S11 and their captions are provided in excel datasheets in the zip file. Table S4-S6 have a shared data description that is applicable to the whole workbook.

Supplementary Methods

#### Main drought screen

##### Experimental design and growth conditions

We grew plants in SC-10 Cone-tainersTM (3.81 cm diameter, 164 ml) in RL98 racks placed into FLOWTS flow trays (Stuewe and Sons, Tangent, Oregon, USA) in a walk-in multi-tier Conviron MTPS growth chamber (Conviron Ltd., Winnipeg, Canada). To mimic more natural growing conditions, we programmed the temperature and photoperiod in the chamber based on that of the *Lip-0* ecotype that originated from southern Poland, representing a common field environment, following Lorts and Lasky (2020) (Table S2). Each tray represented either a drought or well-watered treatment, and seven replicates per mutant were randomly assigned positions for each treatment. Seeds were stratified in DI water for 4-5 days prior to planting.

All pots were well-watered by maintaining 2.5-5 cm of water at the bottom of each tray prior to drought treatments. Drought treatments began 19 days after planting when all seedlings had at least the first true leaves expanded. We removed the bottom water in the drought trays while maintaining a constant 2.5 cm of water in the well-watered trays. To prevent the topsoil from drying, well-watered pots were top-watered every other day with 2.5 ml of water. The soil in drought treatment plants was allowed to dry for five weeks after the bottom water was removed, then we top-watered each pot in both well-watered and drought treatments with 12ml 2x strength miracle grow solution every other day for all trays to maintain adequate nutrition. Trays were rotated and moved to a different bench in the growth chamber every other week. Percent volumetric water content was monitored every 30 minutes throughout the experiment using 3 drought-treatment pots and 2 well-watered pots containing *Columbia-0* (*Col*) and 5TE probes (METER Group, Inc., Pullman WA, USA).

##### Plant harvest and phenotyping at maturity

The fecundity of Arabidopsis plants that survived to reproduce was measured when all siliques reached maturity (dried and brown) and the rosette leaves had senesced. For each Arabidopsis plant, we measured inflorescence height and the number of secondary inflorescence branches (inflorescence branches arising from the inflorescence main axis). We also measured the total silique number and silique length of six siliques corresponding to 10, 20, 40, 60, 80, and 90th percentiles of silique positions along the inflorescence to represent silique length across the entire inflorescence (e.g. the 10th percentile silique was higher up the inflorescence than 10% of all siliques). After siliques were measured, we dried the mature inflorescence and rosette at 60 °C for 24 hours, weighed them separately, and then added their weights to get aboveground dry biomass. Finally, the rosette tissue was ground and used for δ^13^C and δ^15^N isotope analysis at the UC Davis Stable Isotope Facility.

#### Follow-up drought and freezing experiments

##### Intermediate drought experiment on *wrky38* and *lsd1*

We applied a constant temperature setting (20/14℃, 12h/12h) throughout this experiment and an intermediate drought by controlling the frequency of pot saturation. Specifically, we bottom-watered pots every 3 days in the well-watered (WW) treatment while reducing the watering frequency for pots under intermediate drought (ID) treatment to every 6 days starting Day 25, when the drought treatment started. At this time point, most plants had 10 leaves, which corresponds to the adult vegetative phase in *Col* (Lawrence-Paul et al., 2023). We grew 12 replicates of *Col*, *wrky38*, and *lsd1* each for WW and ID treatments. On Day 40, we randomly picked 5-6 pots per genotype per treatment for destructive measurements and kept 6 pots per genotype for post-harvest trait assessments. Rosettes were first cut and weighed for fresh weight, and then each of them was put into a petri dish with distilled water under 4 ℃ in the dark overnight to weigh their turgid weight. Next, all the fully expanded leaves were cut, scanned, and then dried separately with the rest of the rosette tissue in an oven at 60 ℃ for 2 weeks before leaf and rosette dry weights were both weighed. The total leaf area of each plant was analyzed by ImageJ, and the Specific Leaf Area (SLA) of each plant was calculated by (Total leaf area/Total leaf dry weight). Relative Water Content (RWC) was calculated by (Rosette fresh weight - Rosette dry weight) / (Rosette turgid weight - Rosette dry weight). We kept the dried materials and sent samples of lines that showed interactions with drought for δ^13^C isotope analysis, an estimate for water use efficiency (WUE), along with leaf C:N and δ^15^N.

Before the first plant flowered, we tracked stomatal conductance (g_sw_) and Fv/Fm for 10 continuous days using a LI-600 porometer and fluorometer (LI-COR Environmental, Lincoln, NE, USA). g_sw_ measurements were taken daily, approximately one hour before lights were turned off. On the watering days, two measurements were taken: one before watering and another approximately 30 minutes after. Fv/Fm measurements were taken once per day, 20 minutes after the lights were turned off. For each measurement, four pots per genotype per treatment were randomly selected. Measurements were taken on the latest fully expanded leaves that were large enough for proper device application.

We recorded the flowering time for each non-destructively sampled plant. When all plants were mature and started senescing, we counted branch numbers and silique numbers. We measured inflorescence length and the lengths of 12 siliques (or all siliques if fewer than 12) from the top, middle, and bottom of the inflorescence to calculate the average silique length. Finally, we calculated the total silique length as the product of silique number and average silique length. Representing reproduction level, silique number, average silique length, and total silique length were considered fitness estimates.

##### Overnight freezing experiment on *wrky38* and *lsd1*

To mimic the natural freezing stress that Arabidopsis could undergo in its growing season, we set the night (12h) temperature in the growth chamber to -2℃ and the day temperature to 10℃ as our freezing treatment, which started on Day 30 when plants were established and had been under cold acclimation at 10/4℃ (12/12h) for 2 weeks. We surrounded all pots with extra soil for insulation to mimic the circumstances in nature, preventing unnaturally cold soil temperatures.

On day 60, we measured diameter as an estimation of growth under freezing. Starting from Day 64, we shifted the temperature back to 20/14℃ until the end of the experiment to allow plants to flower and then recorded flowering time. We grew the plants until senescence, after which we measured aboveground biomass, silique numbers, inflorescence length, and average and total silique lengths as described above. We grew 12 replicates for each genotype, and all of them survived until the end of the experiment.

#### References

### Lawrence-Paul, E.H., Poethig, R.S., Lasky, J.R., 2023. Vegetative phase change causes age-dependent changes in phenotypic plasticity. New Phytol. 240, 613–625. https://doi.org/10.1111/nph.19174

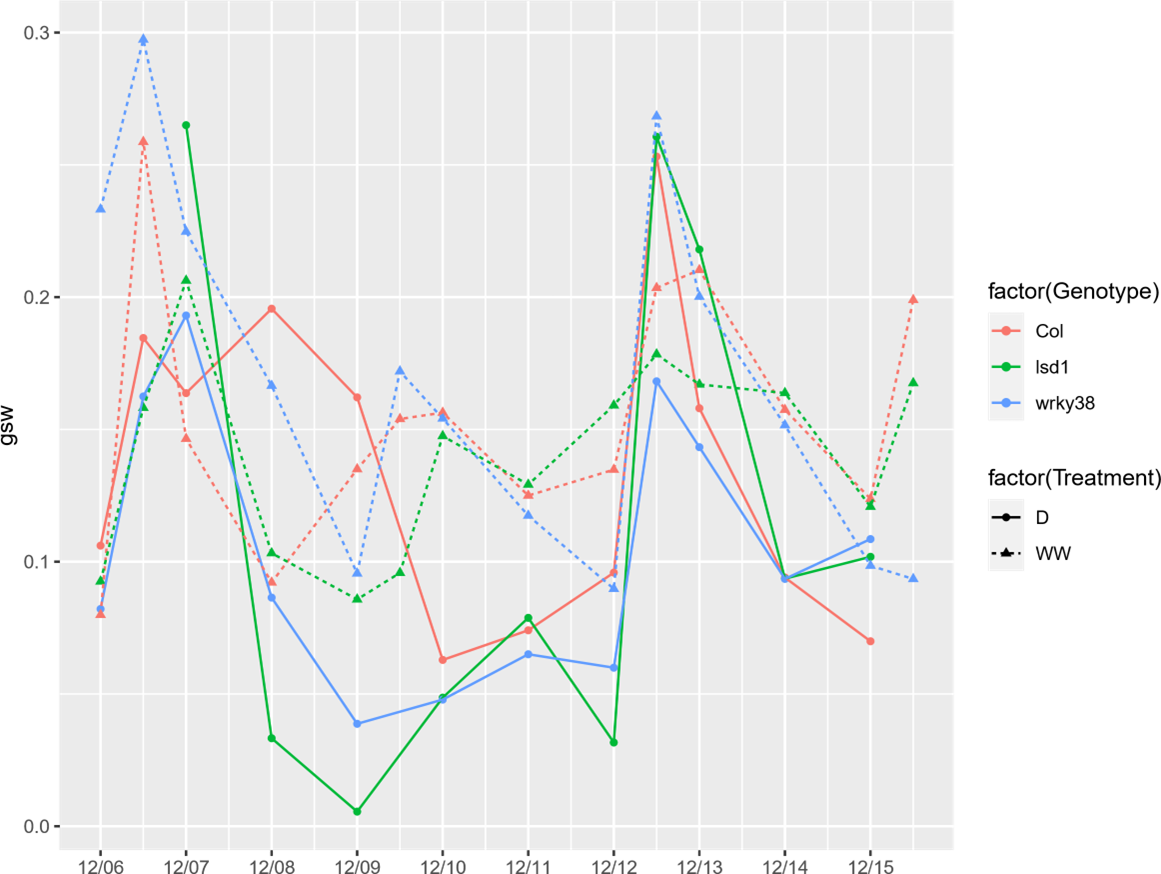

**Figure S1** Stomatal conductance (g_sw_) changes across 10 consecutive days in *Col*, *wrky38*, and *lsd1* measured with LI-COR 600. Measurements were taken at ~1.5 h before lights were off. On the watering day, two g_sw_ measurements were taken right before and ~0.5h after watering. Each point represents mean g_sw_ from 3-4 plants of the same genotype, except that only 2 *lsd1* plants were measured on 12/6/2022. Well-watered (WW) plants were watered on 12/6/2022, 12/9/2022, 12/12/2022, and 12/15/2022. Drought (D) plants were watered on 12/6/2022 and 12/12/2022.

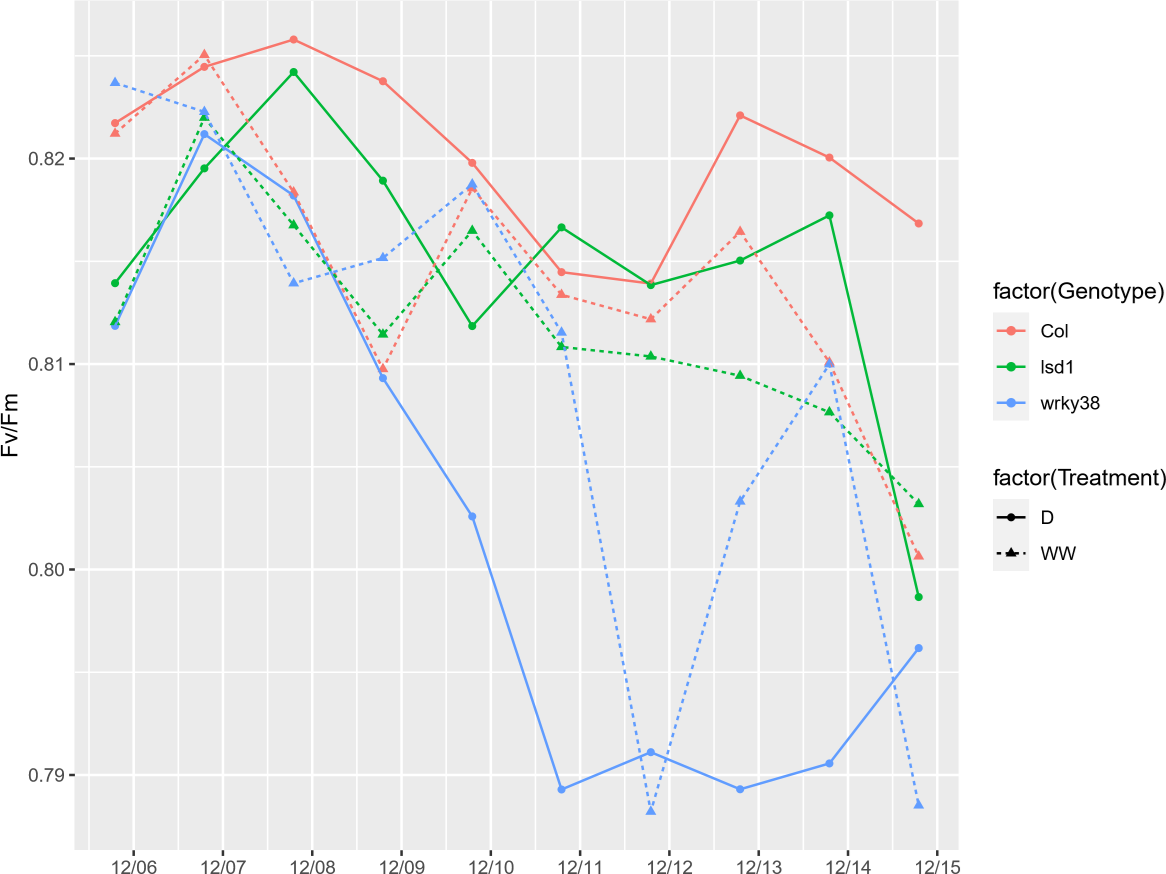

**Figure S2** Fv/Fm changes across 10 consecutive days in *Col*, *wrky38*, and *lsd1* measured with LI-COR 600. Measurements were taken at ~0.5h after lights were off. Each point represents mean Fv/Fm from 3-4 plants of the same genotype, except that only 2 *lsd1* plants were measured on 12/6/2022. Well-watered (WW) plants were watered on 12/6/2022, 12/9/2022, 12/12/2022, and 12/15/2022. Drought (D) plants were watered on 12/6/2022 and 12/12/2022. On the watering day, two Fv/Fm measurements were taken right before and ~0.5h after watering.

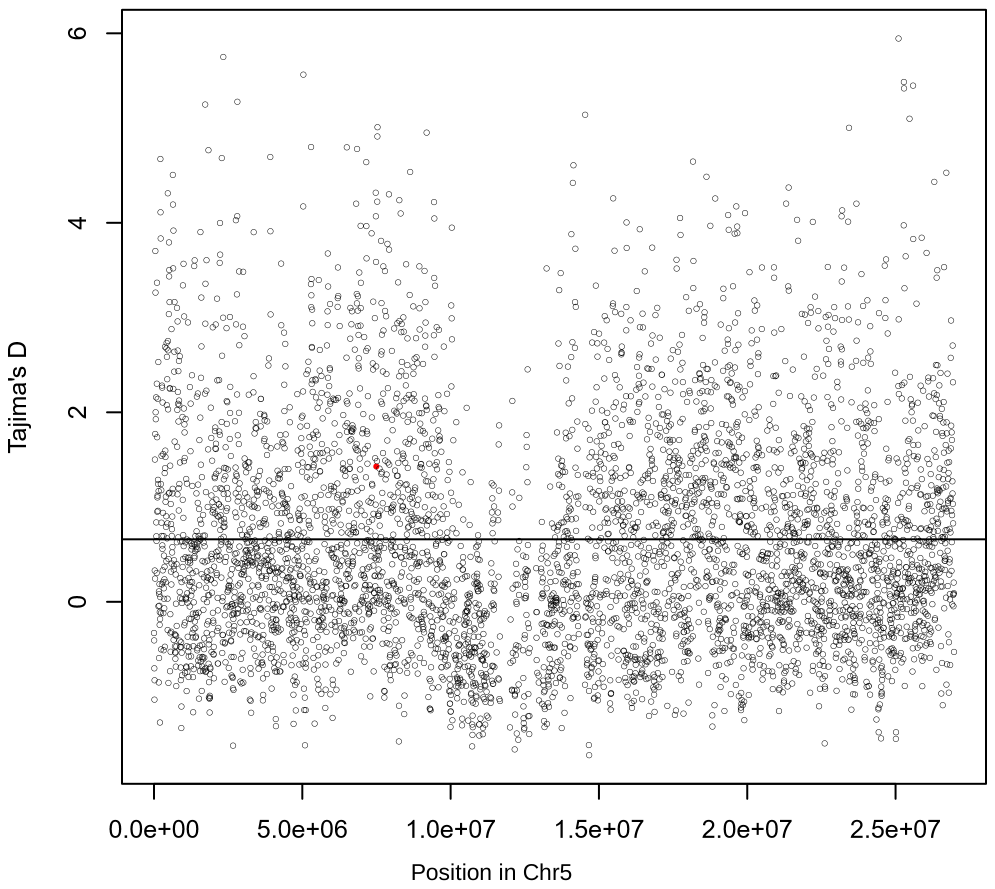

**Figure S3** Tajima's D calculated using 5-kb sliding window across chromosome 5. The black line represents the mean Tajima’s D value across chromosome 5. The red dot represents the window that contains *WRKY38*.

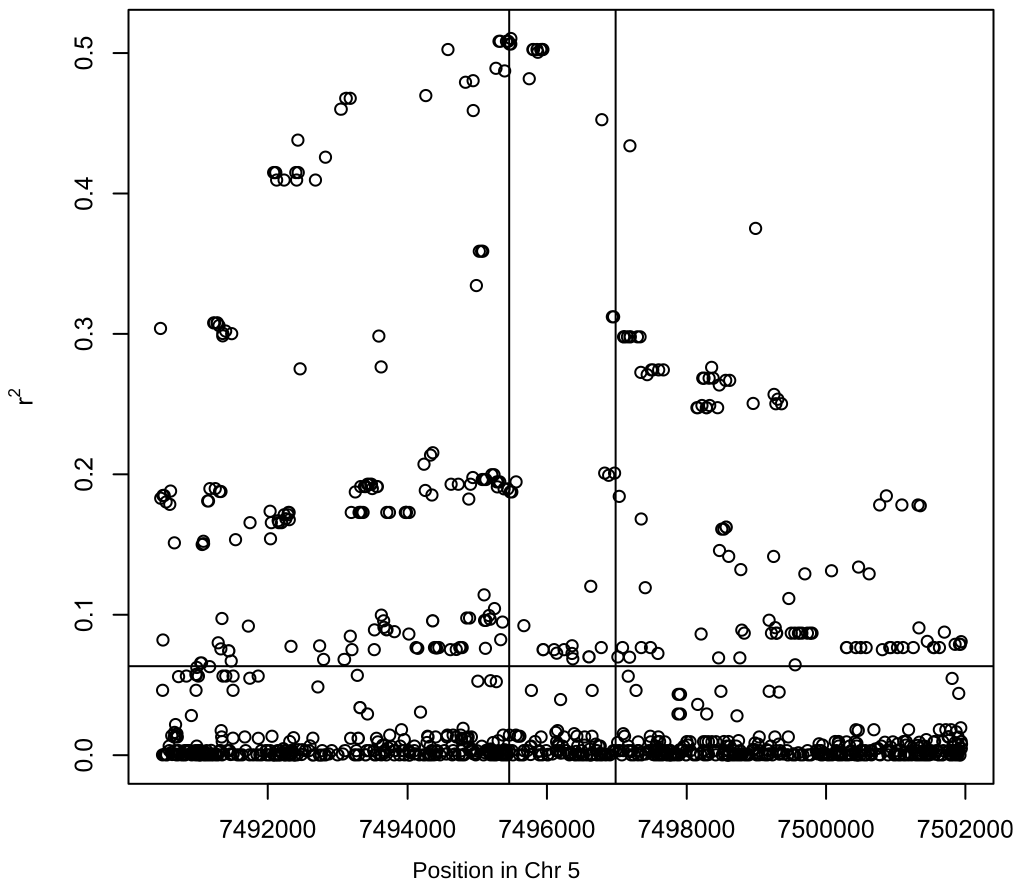

**Figure S4** The square of Spearman’s correlation coefficient (*r^2^*) between Frameshift_7495793_ and each SNP within 10 kb around *WRKY38.* Vertical lines indicate the boundaries of the WRKY38 gene, and the horizontal line represents the mean correlation coefficient within the region.

**Figure S5** SNP heatmap matrix of 1135 accessions from the 1001 Genomes Project within 10 kb of WRKY38. Columns represent SNPs along chromosome 5, rows represent accessions, and white or red squares indicate reference or alternative alleles, respectively. Vertical solid lines mark the boundaries of the WRKY38 gene. Left strips indicate functional variant (FV) types for each accession: teal = frameshift, orange = stop gained, purple = stop lost, pink = intact (functional). Horizontal dashed lines separate accessions based on FV type or position of the FV within the WRKY38 gene. Accessions are ordered by (1) FV type, (2) FV position, and (3) longitude.
