## Supplementary figures and images for "Experimental validation of genome-environment associations in Arabidopsis"

### FigureS1-gsw.pdf

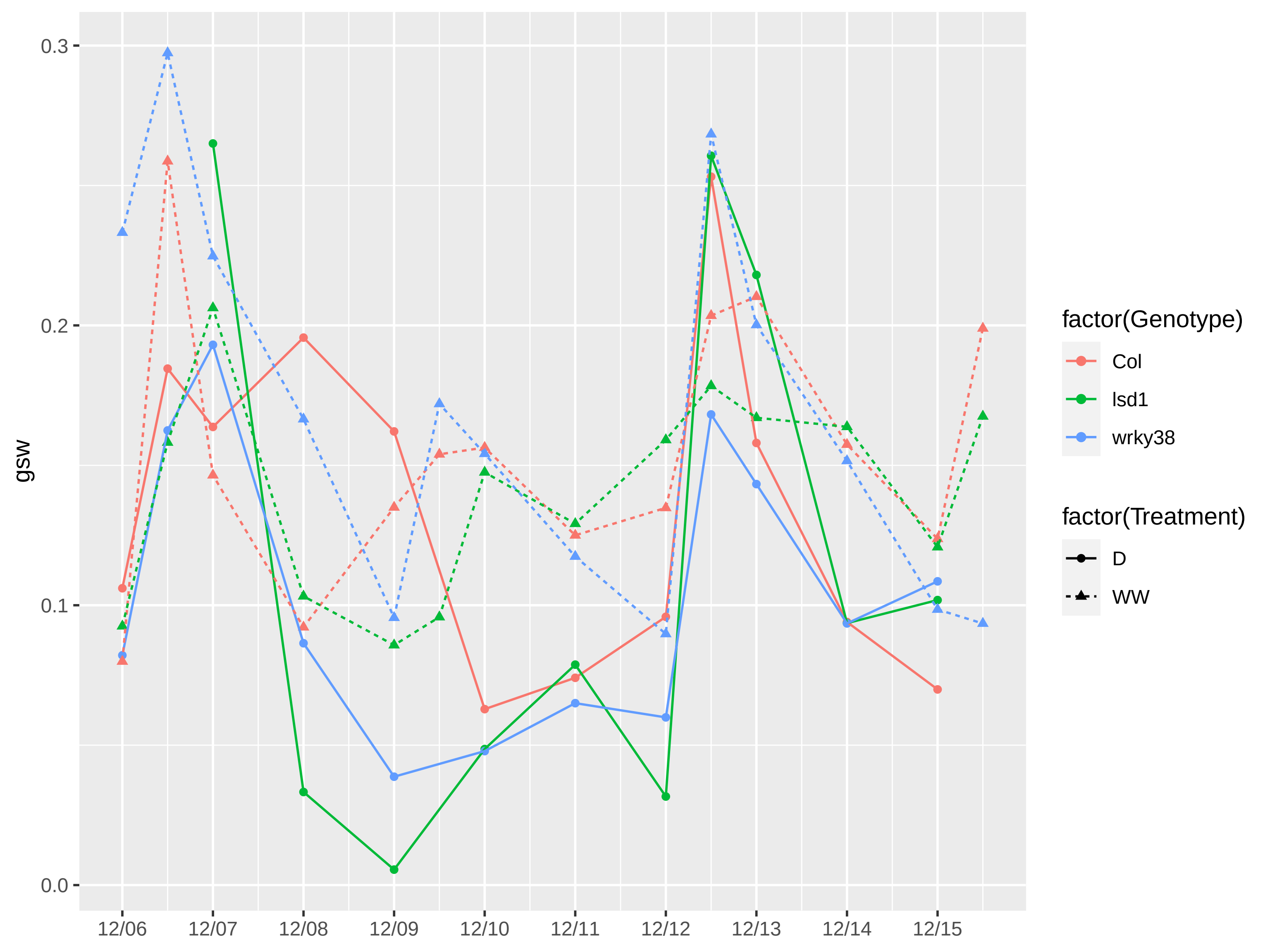

### FigureS2-FvFm.pdf

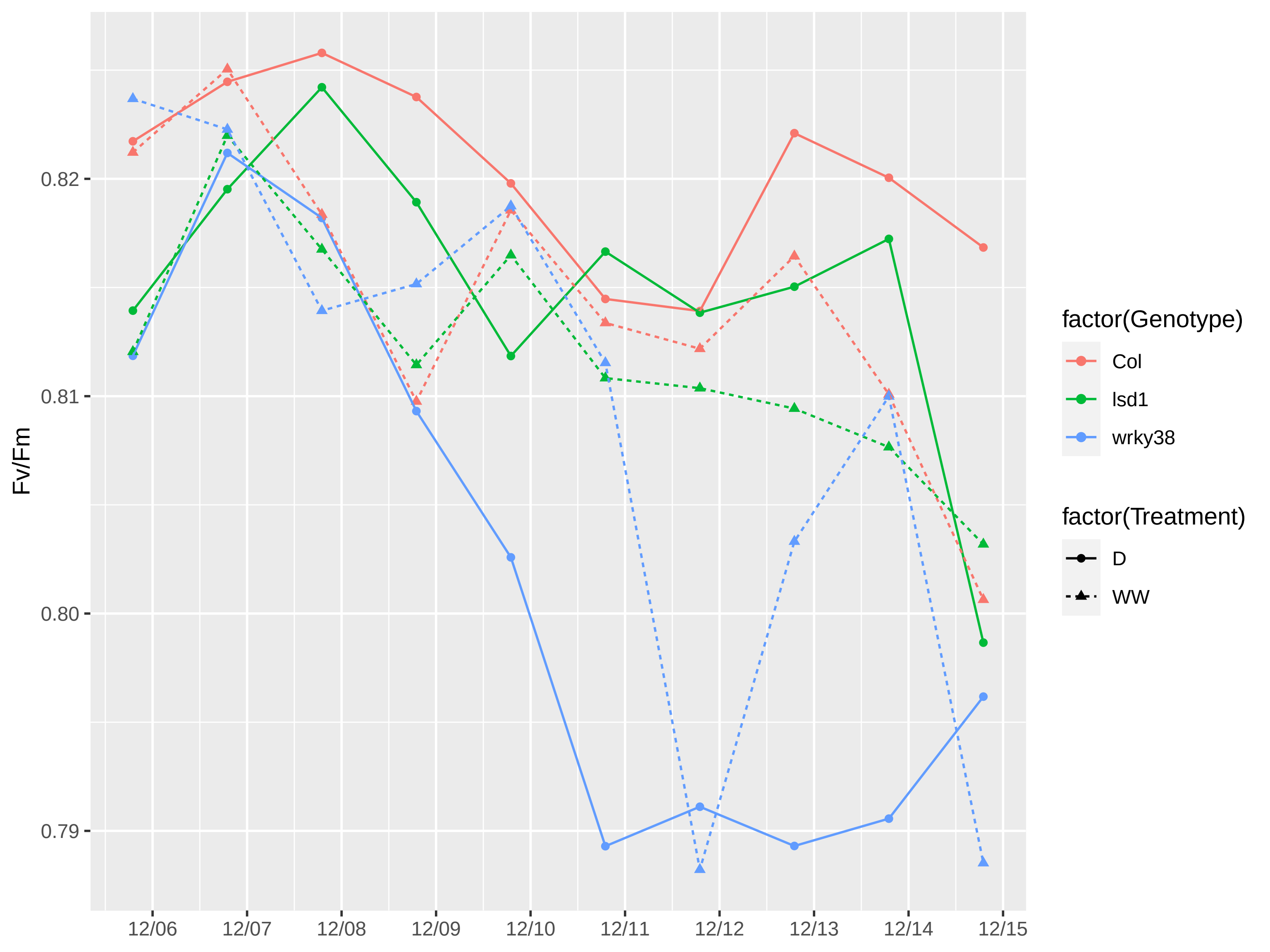

### FigureS3-TajimaD.pdf

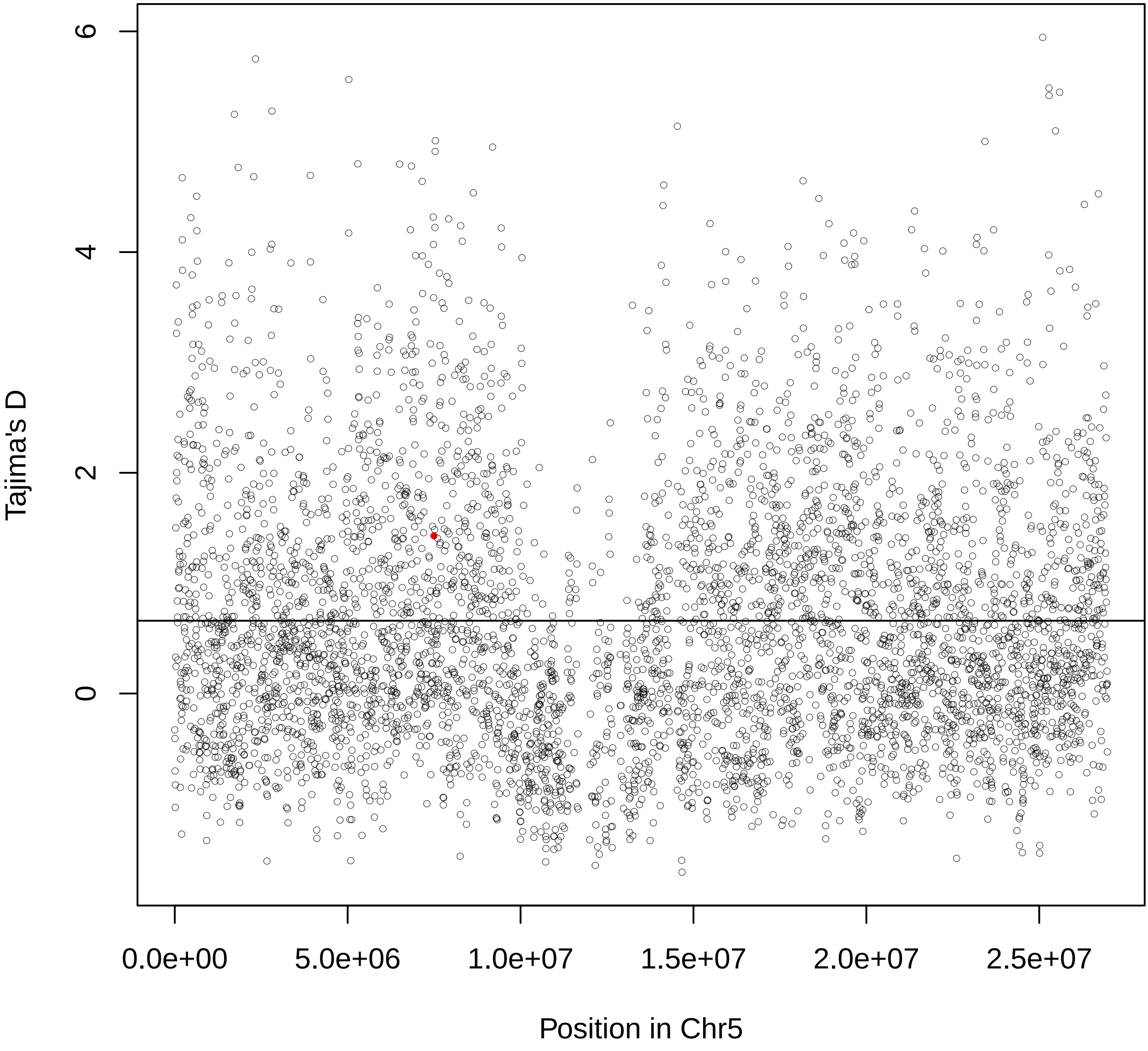

### FigureS4-r2.pdf

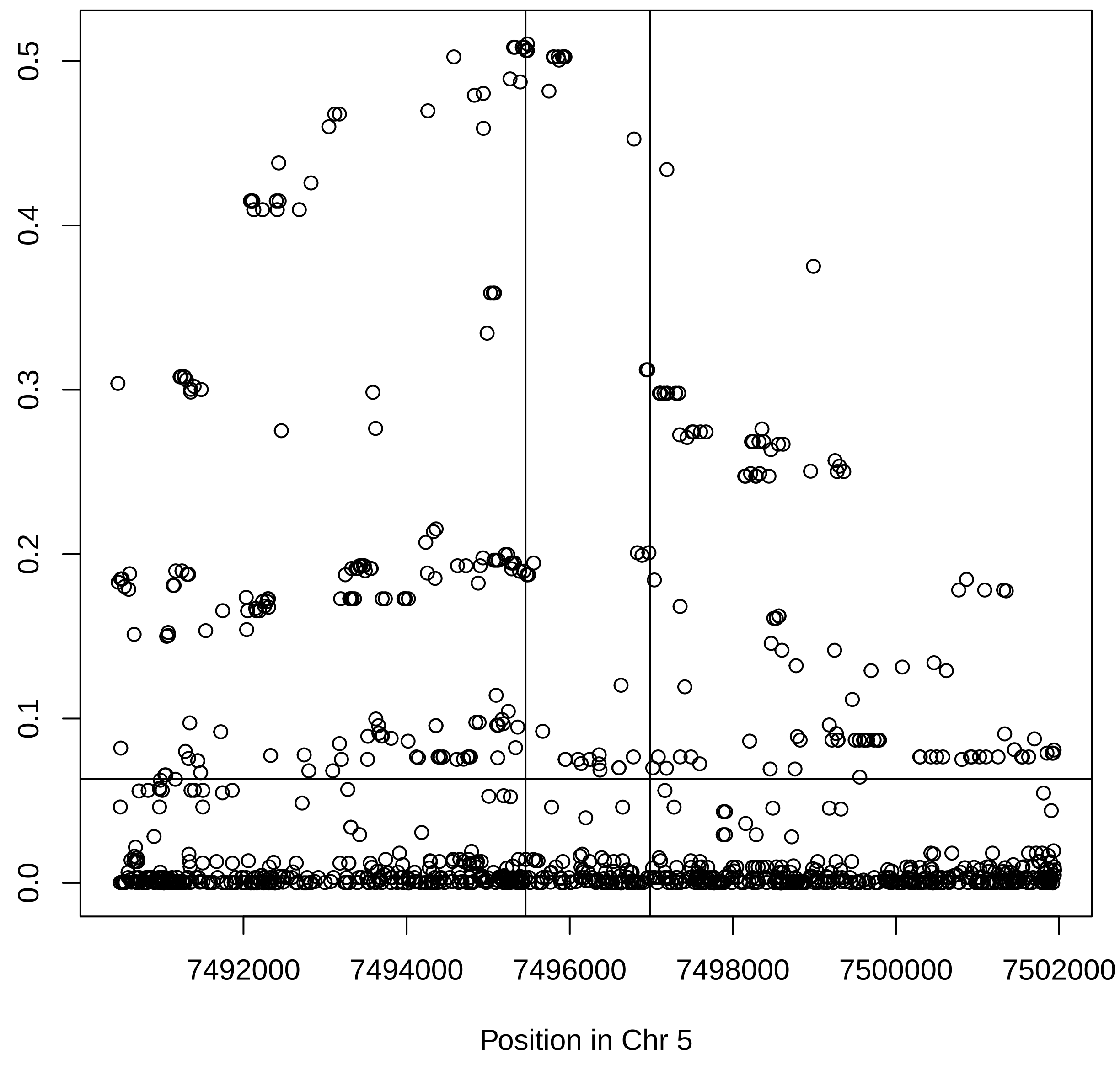

### FigureS5.pdf

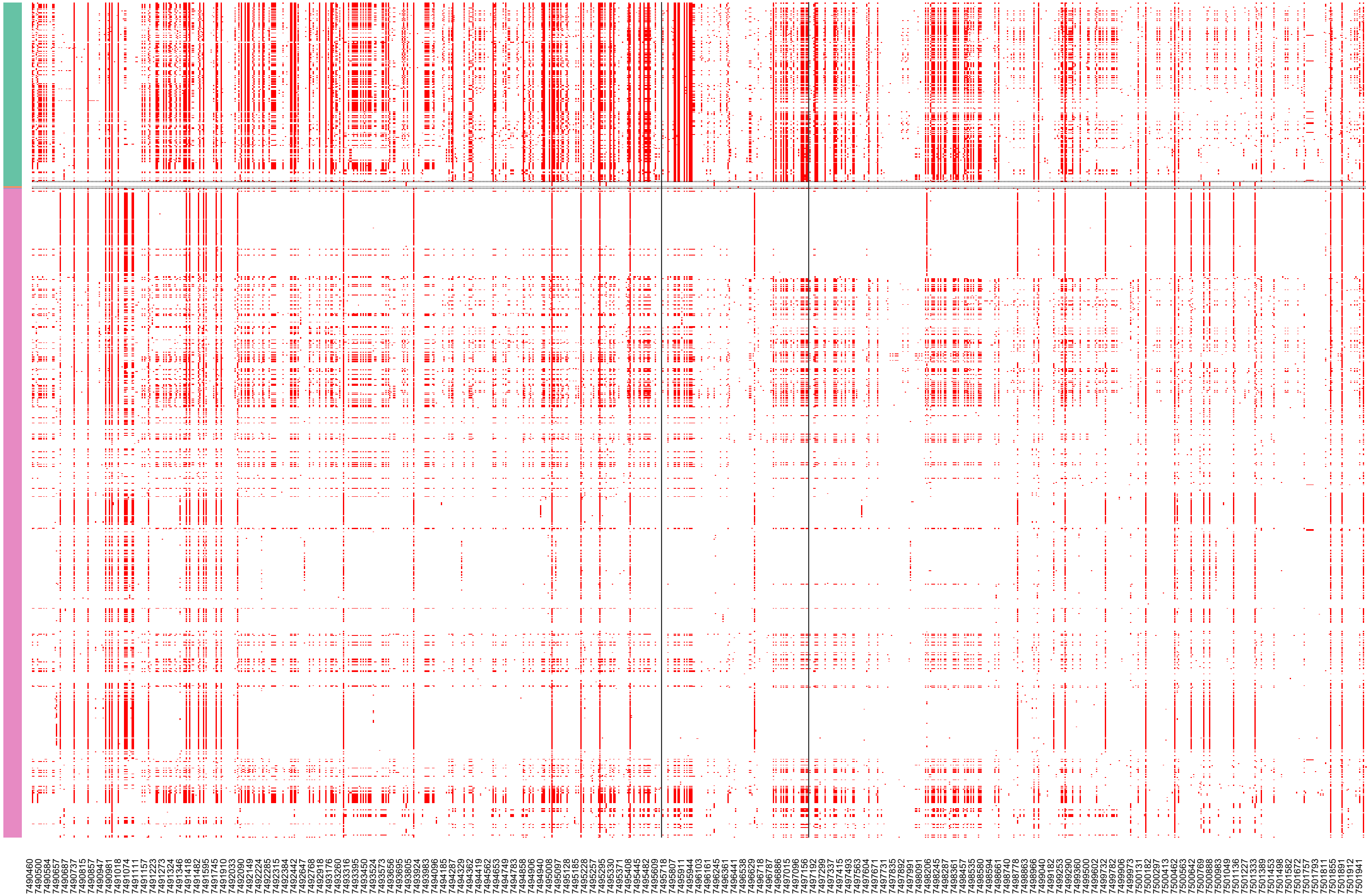
